# Microglia from Friedreich Ataxia patients are intrinsically primed for neuroinflammation

**DOI:** 10.64898/2026.07.16.738960

**Authors:** Ye Man Tang, Rita Lo, Louise Thiry, Michael Fiorini, Sali Farhan, Massimo Pandolfo, Stefano Stifani

## Abstract

Friedreich Ataxia (FRDA) is an autosomal recessive neurodegenerative disorder characterized by progressive loss of cerebellar and proprioceptive neurons that control movement and coordination. In most patients, FRDA is caused by homozygous GAA trinucleotide repeat expansions in the first intron of the *frataxin* (*FXN*) gene, resulting in reduced expression of frataxin, a mitochondrial protein essential for biogenesis of iron-sulfur clusters and mitochondrial function. Although recent therapeutic advances have provided modest clinical benefit, effective disease-modifying treatments remain lacking. Increasing evidence indicates that microglial cell dysfunction contributes to FRDA pathogenesis, highlighting these cells as potential therapeutic targets. However, the molecular mechanisms underlying FXN-deficient microglial dysfunction remain poorly understood. Here, we show that microglia generated from FRDA patient-derived iPSCs exhibit a cell-autonomous pro-inflammatory phenotype in the absence of exogenous inflammatory stimuli. This phenotype is characterized by coordinated activation of immune transcriptional programs, dysregulated secretion of neuroinflammatory proteins, impaired autophagy-lysosomal function, and activation of inflammasomes pathways involving NLRP2 and NLRP3. These findings demonstrate that FXN deficiency is sufficient to induce intrinsic microglial activation and identify molecular pathways that may represent attractive targets for future FRDA therapies.

## Introduction

Friedreich ataxia (FRDA) is an autosomal recessive multisystem disorder whose hallmark neurological manifestation is progressive ataxia. FRDA neuropathology is characterized by the selective vulnerability of specific neuronal populations, including hypoplasia and progressive degeneration of primary proprioceptive neurons in the dorsal root ganglia, followed by degeneration of neurons in the deep cerebellar nuclei and corticospinal tracts. In most patients, FRDA is caused by homozygous GAA trinucleotide repeat expansions in the first intron of the *frataxin* (*FXN*) gene, which induce epigenetic silencing and reduce expression of the mitochondrial protein frataxin (FXN). In a minority of cases, the disease results from a pathogenic loss-of-function *FXN* variant (missense, nonsense, or deletion) in compound heterozygosity with a GAA repeat expansion (Campuzano et al., 1996; Cossée et al., 1999; Reetz et al., 2025). FXN is essential for the biogenesis of iron-sulfur (Fe/S) clusters in the mitochondrial matrix, and its deficiency impairs the activity of multiple Fe/S enzymes, disrupts iron homeostasis and compromises mitochondrial function (Campuzano et al., 1997; Llorens et al., 2019; Reetz et al., 2025; Ristow et al., 2000). Despite recent therapeutic advances, FRDA remains an incurable and progressively disabling disease.

Growing evidence implicates dysfunction of microglia, the resident immune cells of the nervous system, in the pathophysiology of FRDA. Microglia are essential for maintaining neuronal homeostasis and orchestrating neuroinflammatory responses to infection, injury or disease. Under pathological conditions, they acquire an activated state characterized by morphological, transcriptional, and metabolic changes associated with proinflammatory and potentially neurotoxic phenotypes. Activated microglia are present in the dentate nucleus of FRDA patients (Koeppen et al., 2012) and neuroinflammatory changes are evident in the cerebellum and brainstem (Khan et al. 2022). Consistently, FRDA animal models exhibit microglia activation and neuroinflammation (Shen et al., 2016; Della Valle et al., 2023). FXN-deficient microglia from FRDA mouse models display an exaggerated response to inflammatory stimuli together with a shift towards glycolysis and itaconate production, consistent with a pro-inflammatory state (Sciarretta et al., 2024). Moreover, a FRDA mouse model harbouring a human *FXN* transgene with >800 GAA exhibits early cerebellar microglial activation preceding overt disease progression (Vicente-Acosta et al., 2024). Together, these findings suggest that FXN-deficiency drives aberrant microglia activation, creating a pro-inflammatory environment that may contribute to neuronal degeneration. This concept has been directly supported by recent studies using microglia generated from FRDA patient-derived induced pluripotent stem cells (iPSCs), which demonstrated that FXN-deficient microglia are proinflammatory and sufficient to drive neuronal degeneration both in *in vitro* co-culture assays and *in vivo* after implantation into the brains of healthy mice (Pernaci et al., 2025). Overall, these studies establish microgliopathy as an important contributor to FRDA pathogenesis.

However, little is known about the molecular mechanisms underlying the dysregulation of pro-inflammatory mechanisms in FXN-deficient microglia. In this study, we characterized the transcriptome and secretome of microglia generated from FRDA patient-derived iPSCs and matching isogenic controls in the absence of exogenous inflammatory cues. Naïve FXN-deficient microglia are cell autonomously primed for a pro-inflammatory phenotype characterized by dysregulated secretion of numerous neuroinflammatory proteins including several chemokines and cytokines. FXN-deficient microglia display hallmarks suggestive of compromised lysosome biology and autophagic flux, and activation of inflammasome mechanisms involving NLRP2 and NLRP3. These findings identify molecular pathways dysregulated in FXN-deficient microglia that could offer future therapeutic opportunities in FRDA.

## Results

### Cell autonomous dysregulation of pro-inflammatory gene expression in FRDA iPSC-derived microglia

The transcriptional changes underlying the dysregulated activation of human FXN-deficient microglia remain poorly characterized. To address this gap, we performed transcriptomic analysis in FRDA patient iPSC-derived microglia and their corresponding isogenic control cells. Microglia were differentiated from three iPSC lines derived from both female and male FRDA patients. To minimize potential protocol-dependent biases, two different microglia differentiation protocols were used throughout the study (Douvaras et al., 2024; Haenseler et al., 2017). Regardless of donor or differentiation protocol, all microglia preparations consistently displayed typical microglia morphology, and virtually all cells expressed the canonical microglia/macrophage markers P2RY12 and IBA1. Representative images are shown in Figure 1A for microglia derived from the female FRDA iPSC line FRDA_86 (hereafter FRDA_F) and its isogenic control line FRDA_86 GAA homo-del (ISO_F), and from the male FRDA line FRDA_4676 (FRDA_M) and its isogenic control FRDA_4676 GAA homo-del (ISO_M). Bulk RNA sequencing (RNAseq) analysis further confirmed the identity of the differentiated cells, demonstrating robust expression of established microglial marker genes (Butovsky et al., 2014; Hickman et al., 2013), relative to undifferentiated human iPSC lines from publicly available datasets (Fig. 1B). Similar results were obtained with a third FRDA patient-derived iPSC line obtained from a male donor (not shown). We did not detect appreciable differences in the microglia differentiation efficiency or quality among the different FRDA iPSC lines nor between the two differentiation protocols.

**Figure 1.**
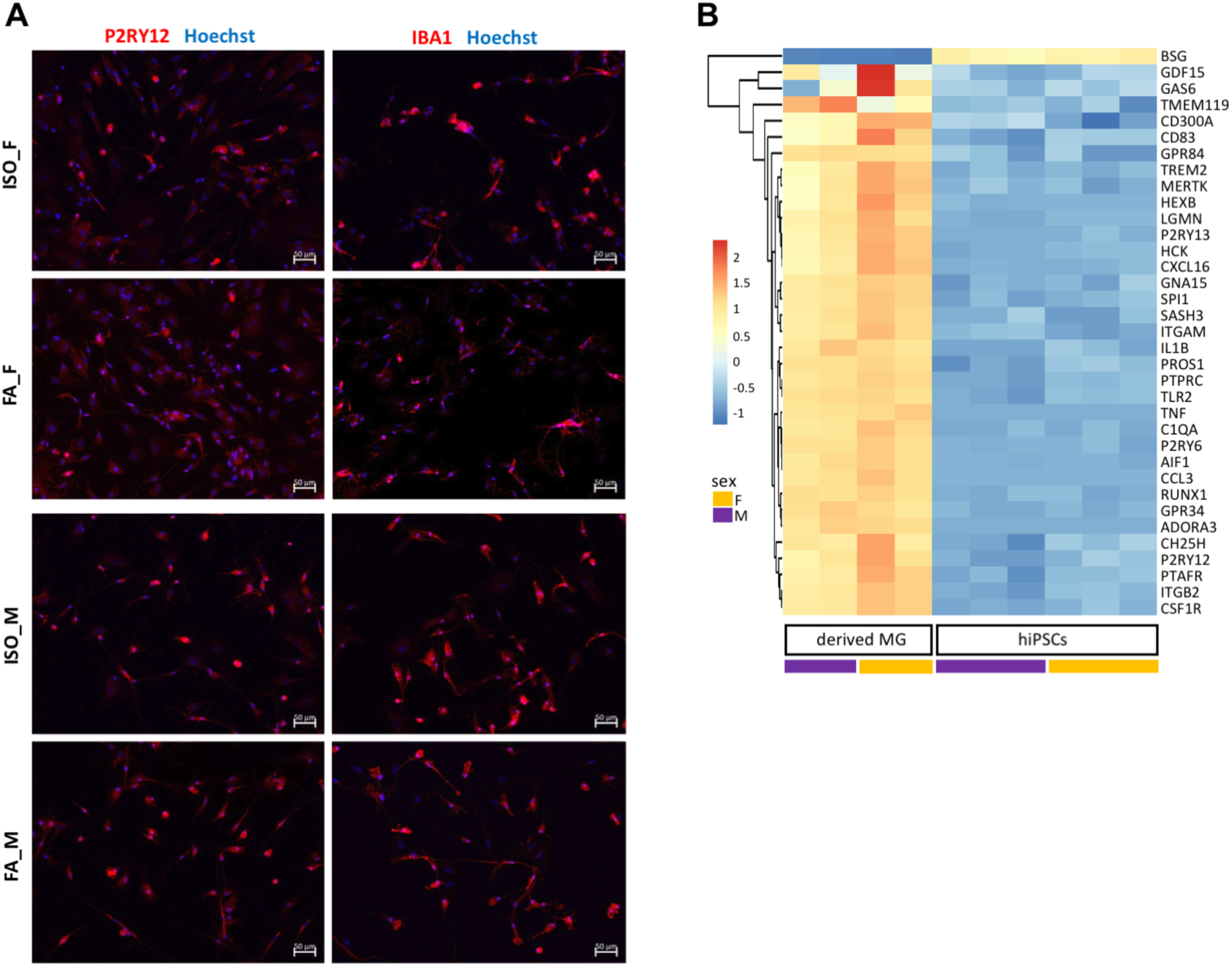
Characterization of iPSC-derived microglia. **A)** Representative images of microglia preparations differentiated from iPSC lines ISO_F, FRDA_F (FA_F), ISO_M, and FRDA_M (F_M), and immunostained with anti-IBA1 or anti-P2RY12 antibodies (red) followed by counterstaining with Hoechst (blue). Scale bar, 50 µm. **B)** Heatmap showing known microglia/macrophage marker gene expression in undifferentiated iPSCs (obtained from publicly available datasets) or microglia preparations differentiated from the two corrected isogenic iPSC lines (ISO_F and ISO_M), used as representative examples. Each column of the heatmap corresponds to one sample (undifferentiated iPSCs (hiPSCs) or isogenic iPSC-derived microglia (derived MG), as indicated at the bottom, and each line corresponds to one gene. In this and succeeding figures depicting heatmaps, subgroups of genes with correlated expression patterns are portrayed by the dendrogram on the left side of each heatmap. Relative levels of gene expression are indicated by a color scale, with cool colors corresponding to low expression and warm colors corresponding to increased/high expression. Hierarchical clustering of rows was applied to the heatmap to group genes with correlated expression patterns.

Microglia derived from both female (FRDA_86) and male (FRDA_4676) iPSC lines were subjected to RNAseq together with isogenic controls. In all experiments (n=8), microglia were maintained under basal conditions, without exposure to lipopolysaccharide (LPS) or any other treatment that would trigger innate immune responses, to characterize the cell-intrinsic transcriptomic changes associated with FXN deficiency. After correction for batch and sex effects, principal component analysis showed clear separation of FRDA and isogenic microglia along one dimension, with samples clustering according to genotype (Fig. 2A). Differential gene expression analysis of iPSC-derived microglia preparations identified 427 up- and 468 down-regulated differentially expressed genes (DEGs) in FRDA microglia relative to isogenic controls (adjusted p-value (false discovery rate; FDR) <0.05, with no fold-change cut-off) (Fig. 2B; entire list of statistically significant DEGs available in Supplemental Information). Among the top 30 upregulated DEGs (Fig. 2C), many encode proteins involved in innate immunity, interferon signaling, and antigen presentation, such as *APOL1*, *CD37*, *NLRP2*, *AQP9, JAK3*, *PADI2*, *APOL3*, *MEG9*, *TMEM176A*, *CD14*, *CALHM2*, *SLC15A4*, *VSIG4, ZNF208, SP140*, *MIR4458HG*, *RYR1*, *PRR5L*, *BST2*, *SVIL-AS1*, *MEG8*, *MEG3* (eg, Cheng et al., 2021; Chiu et al., 2024; Gaudet et al., 2021; Ji et al., 2021; Juliar et al., 2024; Li et al., 2016; Meng et al., 2025; Wang et al., 2013; Xu et al., 2024a; Ye et al., 2023; Zahl et al., 2023; Zhang et al., 2024).

**Figure 2.**
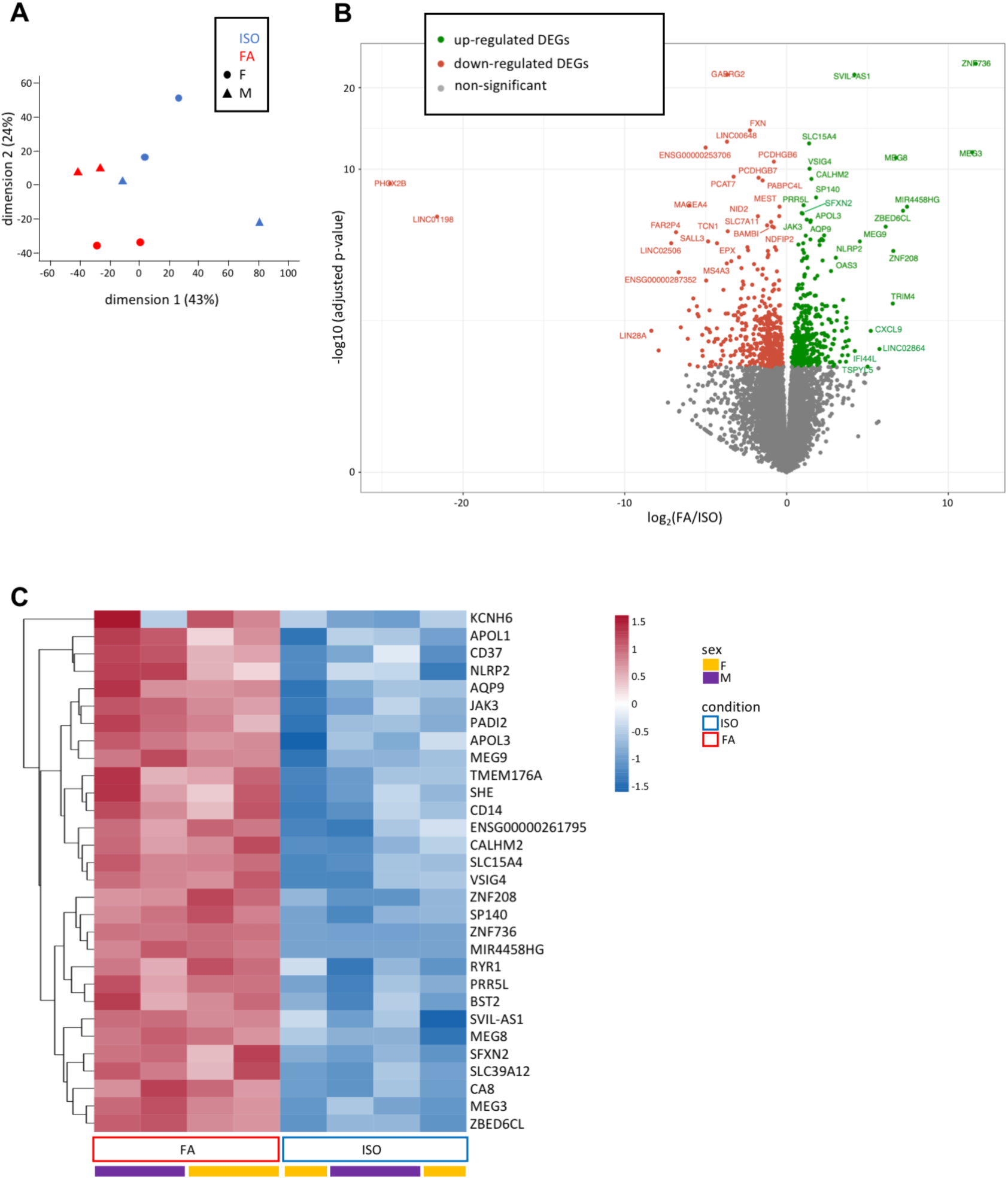
Differentially expressed genes in FRDA *versus* isogenic iPSC-derived microglia. **A)** Principal component analysis plot showing the relationship between the transcriptomic profiles of female and male isogenic (ISO) and FRDA (FA) iPSC-derived microglia. Each plotted data point represents one biological replicate. The symbols refer to the sex (triangles correspond to male samples, dot correspond to female samples). Blue data points correspond to corrected isogenic samples; red data points correspond to FRDA samples. The percentages of variance explained for dimensions 1 and 2 are indicated along the axes. **B)** Volcano plot showing the log Fold-Change (X-axis) and -log10 p value (Y-axis), highlighting the DEGs in FRDA compared to isogenic iPSC-derived microglia. Each plotted data point represents one gene: grey data points correspond to genes that are not differentially expressed, red data points correspond to down-regulated DEGs, and green data points correspond to up-regulated DEGs (p-adjusted <0.05, with no Fold Change cut-off). Names of top dysregulated DEGs are indicated. **C)** Heatmap showing the top 30 up-regulated DEGs in FRDA (FA) compared to isogenic (ISO) iPSC-derived microglia. Each column corresponds to one sample (isogenic or FRDA, as indicated at the bottom) and each line corresponds to one gene.

Over-representation analysis of all the up-regulated DEGs in FRDA microglia revealed significant enrichment of Kyoto Encyclopedia of Genes and Genomes (KEGG) terms related to infection and immunity (Fig. 3A). Consistently, the top 25 over-represented Gene Ontology (GO) Biological Process terms amongst upregulated DEGs were dominated by immune- and inflammation-related categories, including ‘immune response’, ‘inflammatory response’, ‘innate immune response’ (Fig. 3B). In contrast, downregulated DEGs were enriched for KEGG pathways involved in cellular signaling, like “PI3K-Akt signaling”, “Hippo signaling”, “Rap1 signaling” (Fig. 3C), while top over-represented GO Biological Process terms included categories like “regulation of signaling”, “regulation of cell communication”, “animal organ development” (Fig. 3D). Gene Set Enrichment Analysis of the complete ranked gene list yielded highly concordant results (data not shown). Taken together, these findings show that FRDA microglia adopt a cell-intrinsic pro-inflammatory transcriptional program.

**Figure 3.**
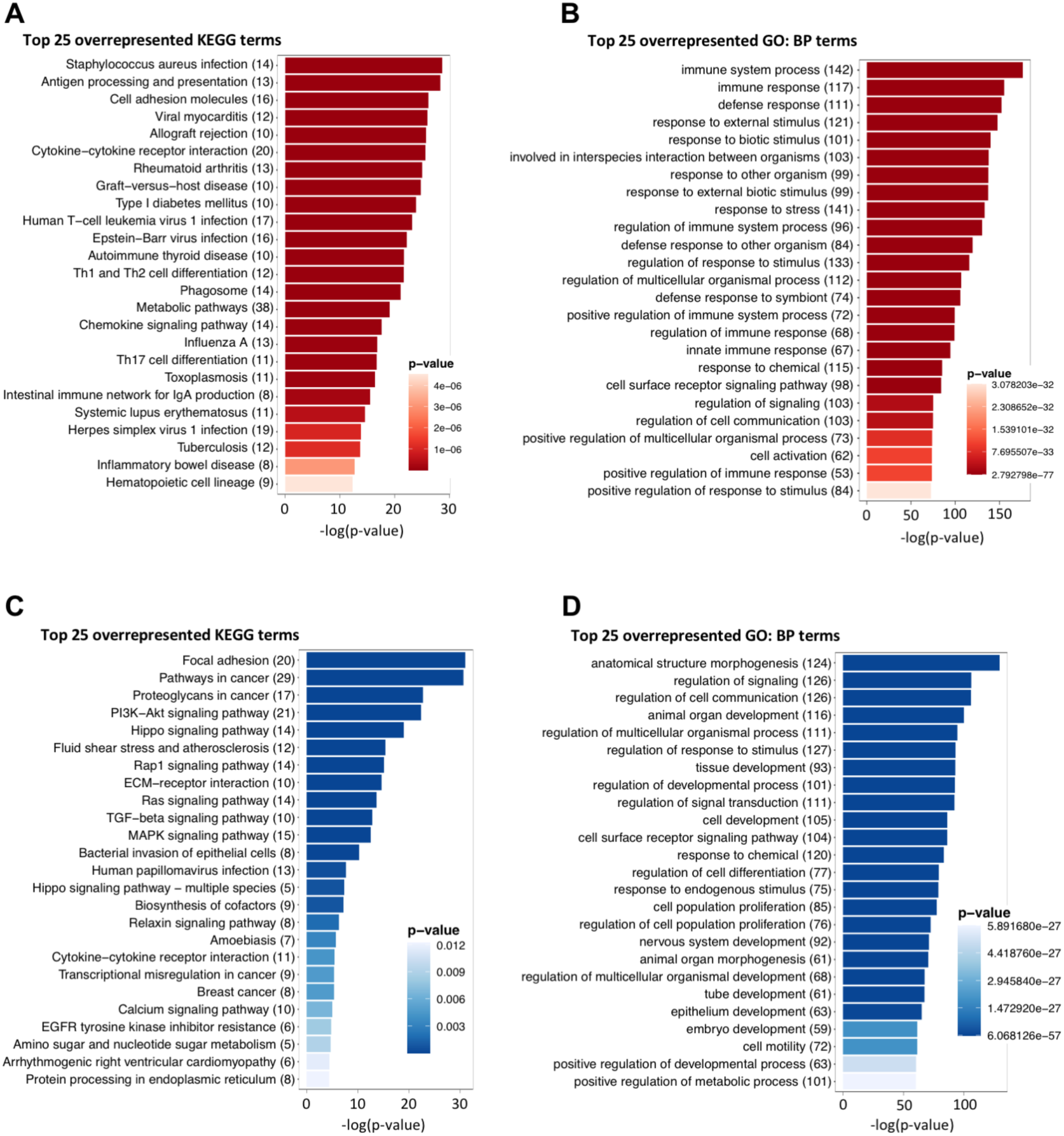
KEGG and GO Biological Process analysis of FRDA microglia DEGs. **A-B)** Bar graphs showing the top 25 over-represented KEGG terms **(A)** and Gene Ontology Biological Processes (GO:BP) terms **(B)** amongst up-regulated DEGs in FRDA microglia. **C-D)** Bar graphs showing the top 25 over-represented KEGG terms (**C**) and GO:BP terms (**D**) amongst down-regulated DEGs in FRDA microglia. The predefined list of significant DEGs was based on FDR-adjusted p-value < 0.05; no fold-change cut-off. For each graph, the number in brackets along the y-axis shows how many DEGs were mapped to the respective term; p values of GO and KEGG terms are indicated with a color-coded scale for each graph, with darker color for smaller values.

### Altered lysosomal homeostasis in FRDA iPSC-derived microglia

Lysosomes play a central role in autophagy/mitophagy by selectively clearing misfolded proteins and damaged organelles. These processes are closely linked to inflammatory signaling, with impaired autophagic flux generally promoting inflammatory responses (Biasizzo and Kopitar-Jerala, 2020; Harris et al., 2017). Consistent with this relationship, previous studies have shown that FRDA iPSC-derived microglia exhibit lysosomal dysfunction, impaired mitochondria-lysosome crosstalk, resulting in impaired mitophagy (Pernaci et al., 2025). In agreement with those findings, GO Cellular Component analysis of upregulated DEGs in FRDA microglia revealed significant enrichment of terms related to intracellular membrane compartments such as “cytoplasmic vesicles”, “organelle membrane”, “lysosome”, “endosome”, suggesting disruption of organelle homeostasis (Fig. 4A). This interpretation was further supported by the presence among the top 30 upregulated DEGs in FRDA microglia of several genes implicated in organelle homeostasis and autophagy regulation, including *SLC15A4* (Lopez-Haber et al, 2022; Rimann et al., 2022), *APOL1* (Blazer et al., 2022), *APOL3* (Ritacco et al., 2026), *RYR1* (Vervliet et al., 2026), *MEG3* (Zang et al, 2022), *CXCL9* (Hu et al., 2025), and *SFXN2* (Chen et al., 2022) (Fig. 2). Conversely, down-regulated DEGs were enriched for terms associated with extracellular components (Fig. 4B). At the protein level, FRDA microglia exhibited decreased expression of lysosomal-associated membrane protein 1 (LAMP1) (Fig. 4C, D), particularly its heavily glycosylated 100-120 kDa forms (Fig. 4F, G). This change was correlated with reduced FXN protein levels (Fig. 4F). Decreased hyperglycosylated LAMP1 has been associated with reduced lysosomal membrane stability and impaired autophagosome–lysosome fusion (Xu et al., 2024b). We also observed that the late endosome marker protein Rab7 trended to decreased levels in FRDA microglia, although this change was not statistically significant under the tested conditions (Fig. 4C, E). Collectively, these findings suggest that, while endosomes may be formed and mainly maintained, the lysosomal compartment is compromised in FRDA microglia.

**Figure 4.**
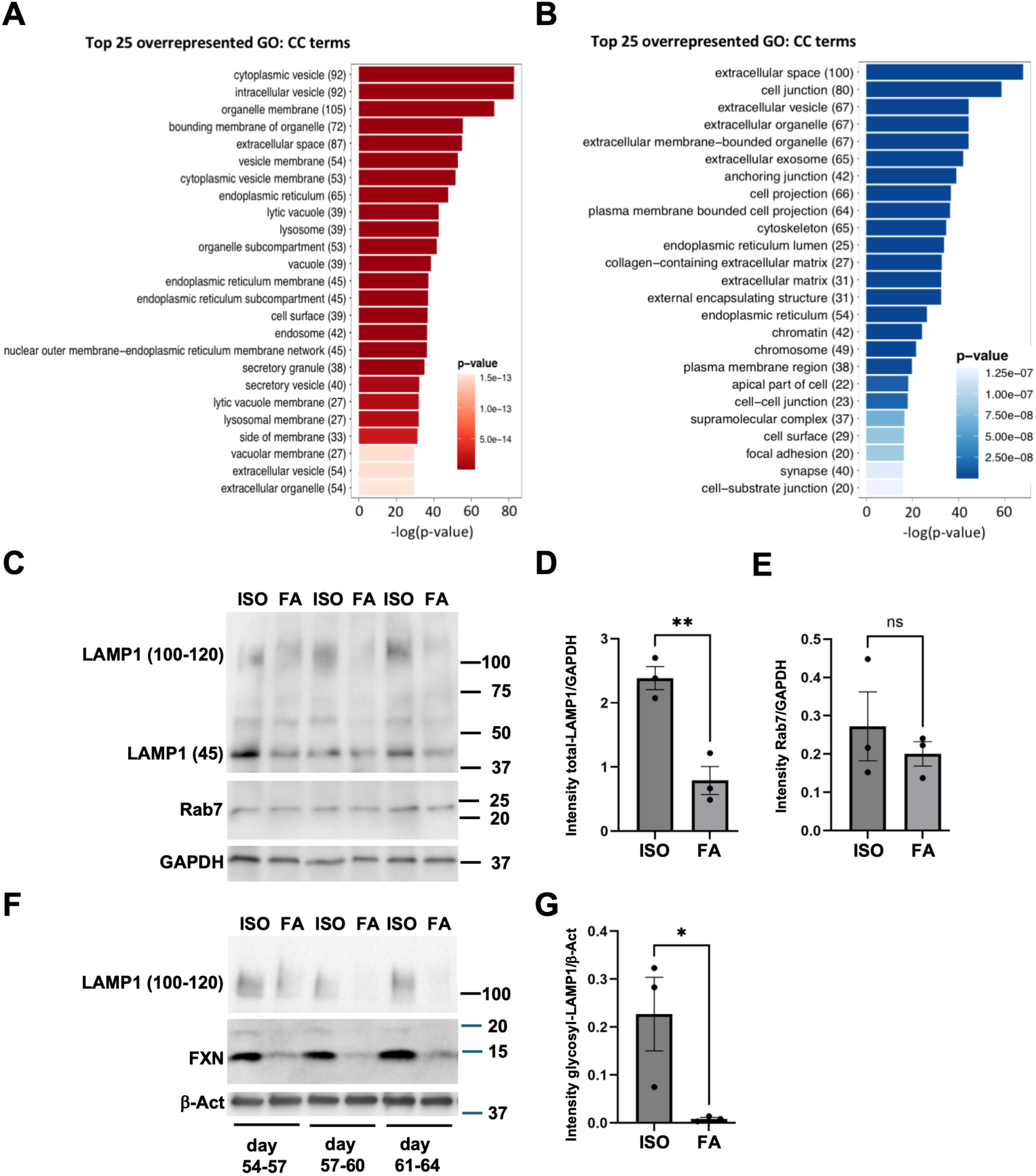
Perturbations of organelle homeostasis in FRDA microglia. **A, B)** Bar graphs showing the top 25 Gene Ontology Cellular Component (GO:CC) terms amongst up- (A) and down- (B) regulated DEGs in both female and male FRDA microglia. The predefined list of significant DEGs was based on FDR-adjusted p-value < 0.05; no fold-change cut-off. **C)** Expression of LAMP1 and Rab7 in microglia derived from iPSC lines ISO_F (ISO) and FRDA_F (FA). Cells were harvested at three increasing stages of *in vitro* differentiation, as indicated below panel F. Basal (∼45 kDa) and hyper-glycosylated (∼100- 120 kDa) LAMP1 forms are visible. **D, E)** Quantification of total LAMP1 and Rab7 expression normalized to GAPDH (ns, not significant). **F)** Expression of hyper-glycosylated LAMP1 and FXN in microglia derived from iPSC lines ISO_F (ISO) and FRDA_F (FA) harvested at the indicated time points. Both mature (∼14 kDa) and intermediate (∼19 kDa) FXN forms are visible. **G)** Quantification of hyper-glycosylated LAMP1 normalized to β-Actin.

### Dysregulated secretion of neuroinflammatory proteins by FRDA iPSC-derived microglia

To determine whether the cell-intrinsic transcriptomic changes observed in FRDA microglia were reflected at the functional level in the microglial secretome, we compared the levels of 285 immune system proteins in the conditioned media from FRDA and isogenic microglia. This analysis identified 57 differentially secreted proteins (DSPs) in the secretome of FRDA microglia, including 43 that were increased (Fig. 5A, C) and 14 that were decreased (Fig. 5B, C) (p value <0.05 and log2FC >0.5). The over-secreted DSPs included several pro-inflammatory members of the CC Chemokine Ligand (CCL) and C-X-C Chemokine Ligand (CCXL) protein families, as well as members of the matrix metalloprotease (MMP) and interleukin (notably IL1-beta (IL1B) and IL6) families, all of which have established roles in neuroinflammation (eg, Becher et al., 2024; Bufi et al., 2025; Li et al., 2025; Sogorb-Esteve et al., 2021; Sun et al., 2025; Wei et al., 2025). Other over-secreted proteins in FRDA microglia included additional pro-inflammatory and neuroinflammatory mediators, like TREM2 (Shi et al., 2025), uPAR/PLAUR (Ma et al., 2026), TLR2 (Heidari et al., 2022), and GZMA (Metkar et al., 2008). These findings indicate that FXN-deficient microglia acquire a pro-inflammatory secretory phenotype.

**Figure 5.**
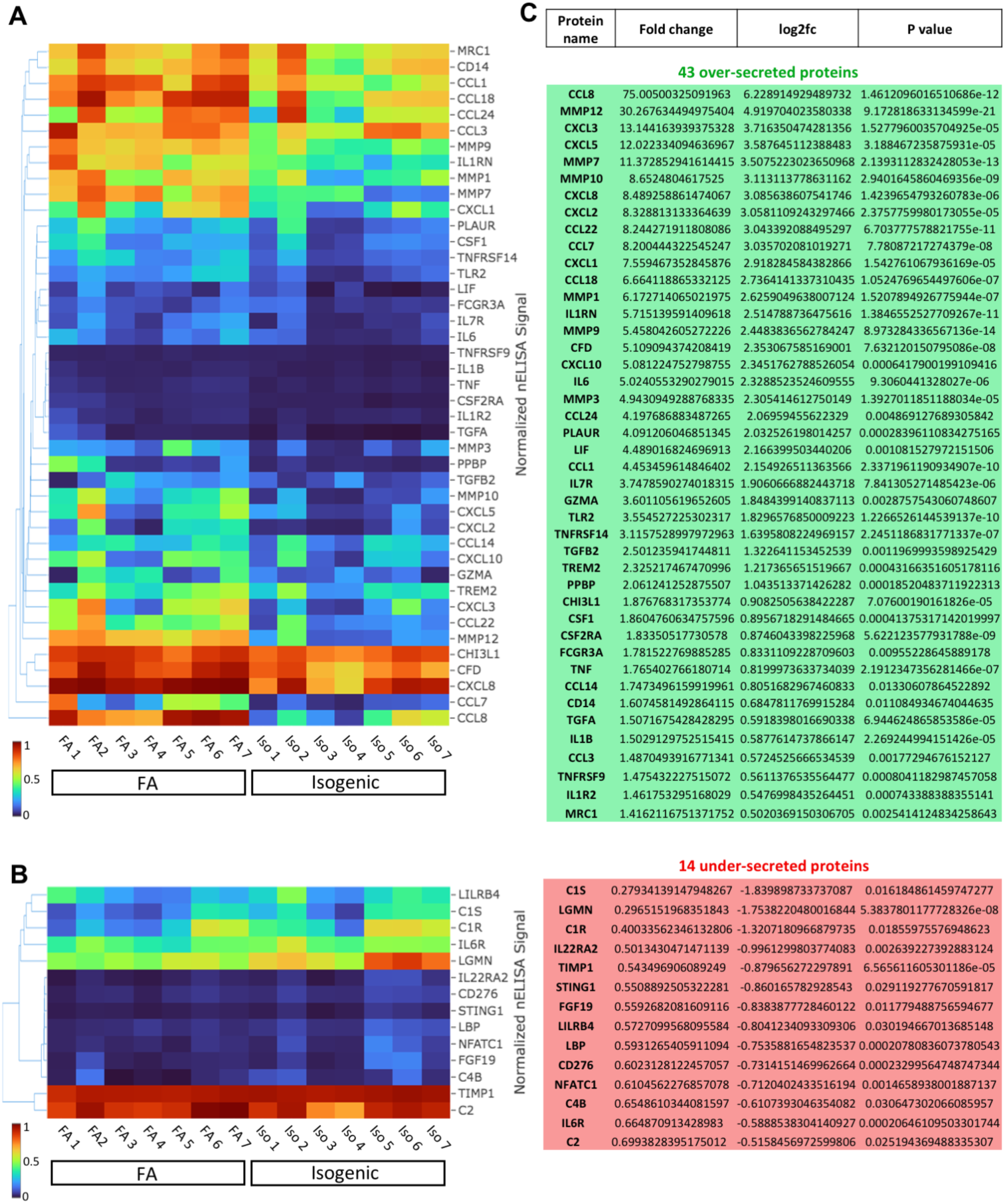
Differentially secreted proteins in FRDA *versus* isogenic microglia. **A, B)** Heatmaps showing the statistically significant (p value <0.05 and log2FC >0.5) differentially secreted proteins in microglia derived from iPSC lines ISO_F (Isogenic) and FRDA_F (FA). Levels of 285 secreted proteins in cell culture supernatants were measured using the nELISA bead-based multiplex immunoassay platform. For each heatmap, columns correspond to separate isogenic or FRDA microglia samples and each line corresponds to one protein. **C)** Table showing the fold change, log2FC, and p values for all the statistically significant over-secreted (43) and under-secreted (14) differentially secreted proteins in FRDA microglia.

Consistent with previous studies showing under-representation of certain complement cascade proteins among differentially expressed proteins in the cerebrospinal fluid of FRDA patients (Imbault et al. 2022), under-secreted proteins in FRDA microglia included the classical complement pathway proteins C1R, C1S and, more marginally, C2 and C4B. In contrast, CFD, a component of the alternative complement cascade, was over-secreted. Collectively, these findings demonstrate broad dysregulation of immune signaling in the FRDA microglial secretome and further support the conclusion that FXN deficiency primes microglia toward a neuroinflammatory state.

### Activation of NLRP2 and NLRP3 inflammasomes in FA microglia

Activation of pro-inflammatory microglia is frequently associated with inflammasome signaling mediated by members of the Nucleotide-Binding-Oligomerization Domain (NOD)- and Leucine-Rich Repeat (LRR)-containing (NLR) family of proteins, particularly NLR Pyrin-Domain-Containing (NLRP) 2 and -3 (Ducza and Gaál, 2024; Lu et al., 2026). A variety of ‘danger signals’, including potentially harmful microbial agents and signs of tissue damage, increase expression of *NLRP2* and *NLRP3*, ‘priming’ the activation of inflammasome responses, which is then completed by assembly of multiprotein complexes that recruit and activate caspase-1 (CASP1). Activated CASP1 processes pro-inflammatory molecules, including but not limited to interleukin-1beta (IL1B), IL18, and IL37 (Ducza and Gaál, 2024; Fu and Wu, 2023; Paik et al., 2025; Sharma and Kanneganti, 2021).

*NLRP2* was among the most strongly upregulated DEGs in FRDA microglia (log2FoldChange, 4.53; p value, 1.529E-08) (Fig. 2). Consistent with transcriptomic data, levels of the full-length NLRP2 protein (∼120 kDa), detected using an antibody directed against the N-terminal pyrin domain and part of the central NACHT domain, were significantly increased in FRDA microglia at several stages of *in vitro* maturation (Fig. 6A, C). We also detected faster-migrating NLRP2 forms of ∼80-90 kDa, possibly corresponding to C-terminally truncated species. NLRP3 protein expression was similarly increased in FRDA microglia (Fig. 6B, D). Consistent with these observations, several top upregulated DEGs in FRDA microglia, such as *AQP9*, *SLC15A4*, *MEG3, PADI2,* and *APOL1* (Fig. 2), are known to promote/mediate NLRP3 inflammasome activation (Guzmán-Guzmán et al., 2021; Lopez-Haber et al., 2022; Meng et al., 2021; Mishra et al., 2019; Wu et al., 2021; Zhu et al., 2023). In contrast, expression of the anti-inflammatory NLR family member *NLRC3*, a negative regulator of both *NLRP2* and *NLRP3* (Shiau et al., 2013; Zhang et al., 2014), was decreased in FRDA microglia (log2FoldChange, -1.721; p value, 0.0405). Together, these observations indicate that FXN deficiency primes NLRP2/NLRP3 inflammasome signaling in human microglia.

**Figure 6.**
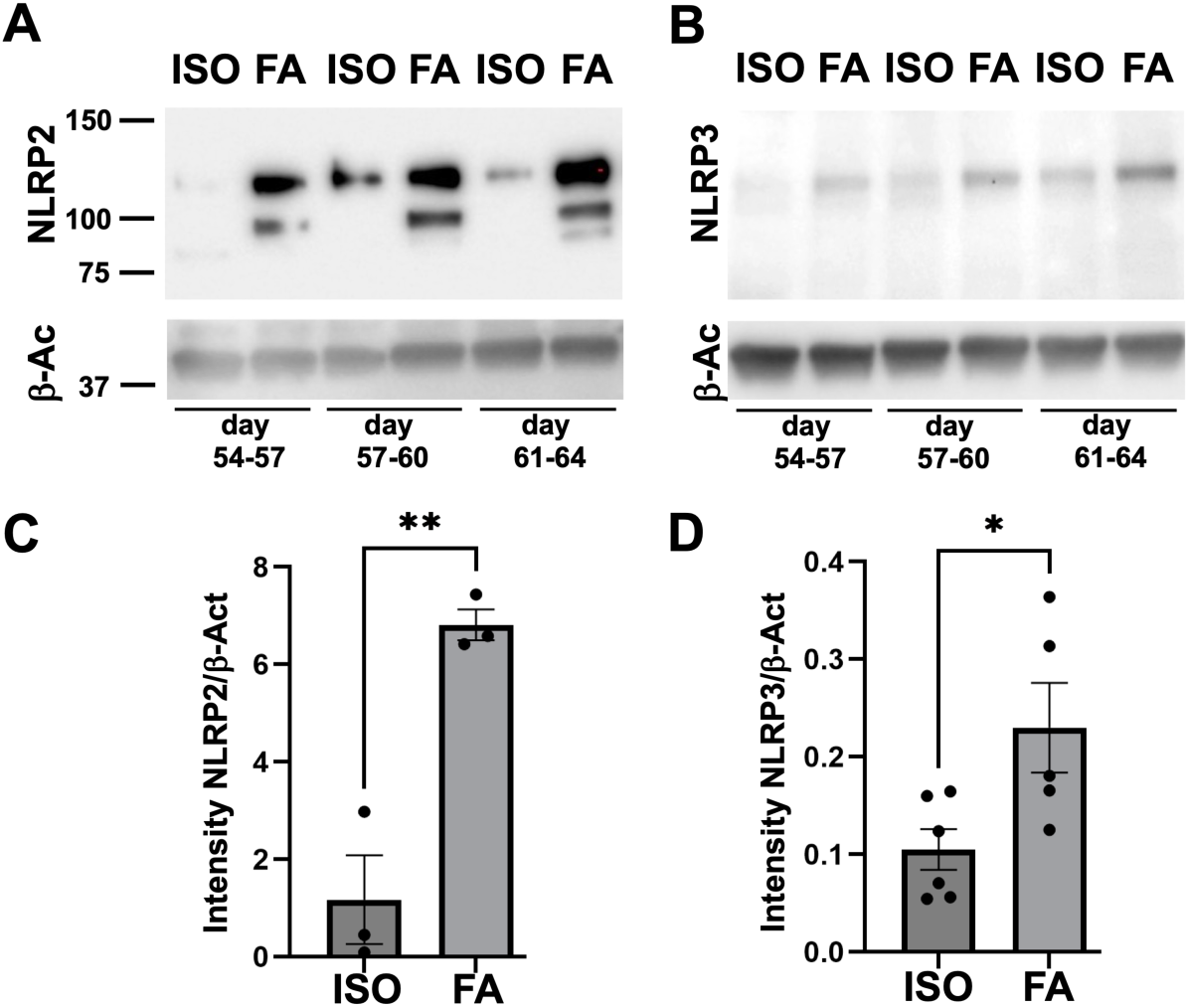
Analysis of NLRP2 and NLRP3 expression in FRDA microglia. **A, B)** Western blotting analysis of NLRP2 (A) and NLRP3 (B) expression in in microglia derived from iPSC lines ISO_F (ISO) and FRDA_F (FA). Microglia were collected at the indicated days after the start of the differentiation protocol. **C, D)** Quantification of protein expression levels normalized to β-Actin.

Following priming, NLRP2 and NLRP3 inflammasome activation requires assembly of supramolecular complexes through recruitment of the Apoptosis-associated Speck-like protein containing a Caspase-activation and recruitment domain (ASC). ASC subsequently oligomerizes and recruits inactive full-length CASP1 to the NLRP2/3-ASC complexes, promoting CASP1 autocatalytic cleavage into smaller catalytically active forms (Ducza and Gaál, 2024; de Alba, 2019; Vajjhala et al., 2012). To determine whether NLRP2/3 inflammasome activation was enhanced in FRDA microglia, we examined ASC expression and oligomerization by immunoblotting both under standard conditions and following protein cross-linking, a method commonly used to stabilize and detect high-molecular-weight ASC oligomers (Prather et al., 2022; Yu et al., 2023). Under non-cross-linked conditions, western blotting detected a roughly 22 kDa band corresponding to monomeric ASC (’Monom.”) and a ∼32 kDa form, consistent with previously described 30-35 kDa post-translationally modified ASC monomers (Friker et al., 2020; Hara et al., 2013) (Fig. 7A). Monomeric ASC levels were unchanged in FRDA microglia, but the 32 kDa ASC immunoreactive species was modestly increased relative to isogenic microglia (Fig. 7B). Cross-linking preserved these bands (Fig. 7D) and additionally revealed higher-molecular-weight ASC species, including a ∼65 kDa band consistent with ASC trimers (”Trim.”) and oligomeric complexes exceeding 250 kDa (“oligom.”). These high-MW ASC^oligom^ species were significantly increased in FRDA microglia, providing evidence for enhanced ASC oligomerization and inflammasome assembly (Fig. 7D, E).

**Figure 7.**
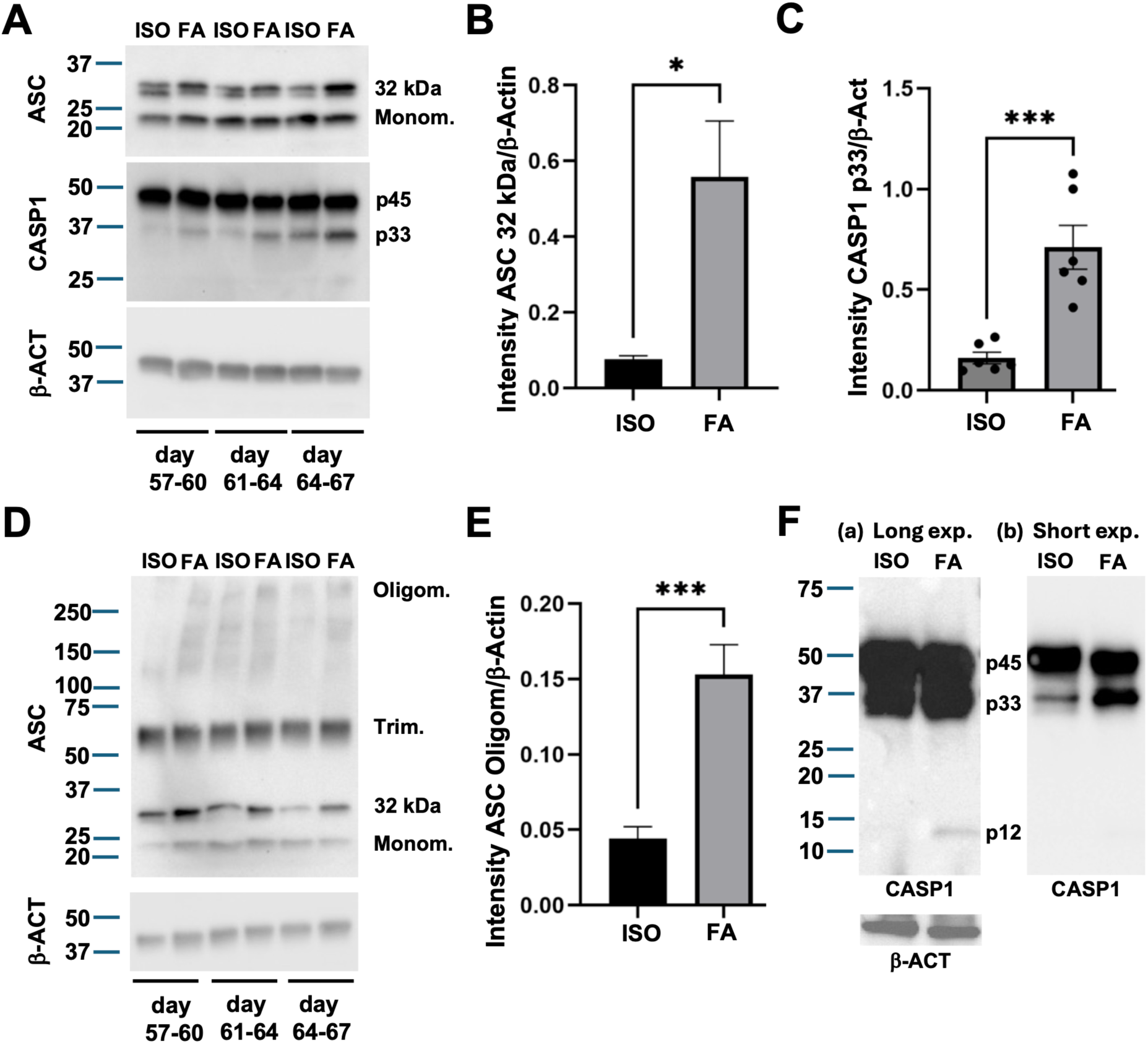
Analysis of ASC and CASP1 expression in FRDA microglia. **A)** Western blotting analysis of the indicated proteins after standard SDS-PAGE. Separate lysates were prepared from microglia derived from iPSC lines ISO_F (ISO) and FRDA_F (FA). Microglia were collected at the indicated days after the start of the differentiation protocol. **B, C)** Quantification of 32-kDa ASC and CASP1 p33 protein expression in FRDA (FA) *versus* isogenic (ISO) microglia normalized to β-Actin. **D)** Western blotting analysis of ASC expression after microglial cell lysates were cross-linked with DTSSP and gels were run under non-reducing conditions. **E)** Quantification of oligomerized ASC expression in FRDA (FA) *versus* isogenic (ISO) microglia normalized to β-Actin. **F)** Expression of CASP1 in microglia derived from iPSC lines ISO_F (ISO) and FRDA_F (FA) and collected 70-73 days after the start of differentiation. (a) Long exposure showing expression of the p45, p33, and p12 forms of CASP1. (b) Short exposure of the same blot.

We next examined CASP1 expression and processing to determine whether the increased expression of NLRP2, NLRP3, and oligomerized ASC was accompanied by CASP1 cleavage and activation. Upon inflammasome activation, pro-CASP1 (45-48 kDa; ‘p45’) initially undergoes autoproteolytic cleavage to generate a transient form of 33-35 kDa (‘p33’, containing the CARD and ‘p20’ (20-22 kDa) domains) and a 10-12 kDa (‘p12’) fragment. The p33:p12 complex is a fully active CASP1 complex that remains bound to the inflammasome supramolecular structure (Boucher et al., 2018). Further cleavage of p33 generates the p20 subunit, which forms an active complex with p12 (Boucher et al., 2018; Makoni and Nichols, 2021). The p20:p12 complex is less stable than p30:p12 dimers and it is thought to contribute to termination of inflammasome signaling (Boucher et al., 2018; Sandstrom and Vance, 2018). Using a CASP1 antibody predicted to react with p45, p33, and p20, we observed immunoreactive species resembling both p45 and p33 in our microglia preparations at approximately two months after the start of the differentiation protocol, but we did not detect p20 (Fig. 7A). Whereas p45 levels were not detectably different, p33 was significantly increased in FRDA *versus* isogenic microglia (Fig. 7C). A different CASP1 antibody, predicted to react with p45, p33, p20 and p12, also did not detect p20 but revealed the p12 form in FRDA microglia, even though this required longer blot exposures than needed to visualize p45 and p33 (Fig. 7F). These results show that naïve FXN-deficient microglia have increased levels of p33:p12 CASP1 dimers, suggestive of augmented CASP1 activation. Overall, the observed increase in NLRP2, NLRP3, oligomerized ASC, and active CASP1 dimers is strongly supportive of a cell-intrinsic activation of inflammasome pathways in naïve FRDA microglia, without exogenous cues.

## Discussion

FRDA is a neurodegenerative disease characterized by progressive loss of vulnerable neuronal populations, particularly cerebellar and proprioceptive neurons involved in motor coordination. Although omaveloxolone provides modest clinical benefit, effective disease-modifying therapies are lacking. Emerging evidence suggests that, as in other neurodegenerative disorders, neuroinflammation contributes to FRDA pathogenesis. Building on previous studies of FRDA *post-mortem* tissues (Khan et al. 2022; Koeppen et al., 2012), FRDA animal models (Della Valle et al., 2023; Sciarretta et al., 2024; Shen et al., 2016; Vicente-Acosta et al., 2024), and FXN-deficient human iPSC-derived microglia (Pernaci et al., 2025), the present study sought to define the molecular basis of the cell autonomous phenotype of FXN-deficient microglia and its potential contribution to disease pathogenesis. To this end, we analyzed naïve human microglia generated from FRDA patients iPSCs under basal conditions, without exogenous inflammatory stimulation. This model provides a unique opportunity to examine the direct consequences of FXN deficiency in human microglia, independently of interactions with neurons or other glial cells. This distinction is important because human and murine microglia differ substantially in their transcriptional profiles, particularly for genes implicated in neurodegenerative disease (Galatro et al., 2017; Healy et al., 2020; Masuda et al., 2019), and because animal models cannot readily distinguish cell-autonomous from non-cell-autonomous mechanisms.

Our findings demonstrate that naïve human FXN-deficient microglia are intrinsically primed toward a pro-inflammatory phenotype characterized by coordinated activation of innate immune transcriptional programs, over-secretion of numerous neuroinflammatory mediators, disruption of lysosomal homeostasis and autophagic flux, and activation of inflammasome mechanisms involving NLRP2 and NLRP3. These observations are consistent with both the pro-inflammatory phenotype of FRDA microglia and previous demonstration that these cells have altered mitochondrial homeostasis, including increased reactive oxygen species (ROS) production, both established triggers of NLRP3 inflammasome activation (Dominic et al., 2022; Liu et al., 2018). Although NLRP3 has been extensively studied in microglia, considerably less is known about NLRP2, whose functions have been investigated primarily in astrocytes, where it is involved in ATP-dependent inflammasome activation and in conditions mimicking ischemic stroke (Cheon et al., 2018; Minkiewicz et al., 2013). Knockdown of *NLRP2* in human astrocytes significantly decreases CASP1 processing in response to ATP, suggesting that the NLRP2 inflammasome is an important component of astrocyte-mediated pro-inflammatory responses mediated by CASP1 (Minkiewicz et al., 2013). NLRP3 is also upregulated by ATP in astrocytes (Albalawi et al., 2017), and astrocytes exposed to Aβ-calcium sensing receptor complexes respond by upregulating the expression of both NLRP2 and NLRP3 (Chiarini et al., 2025).

The similar time course of NLRP2 and NLRP3 upregulation observed in the present study suggests that these inflammasomes may play redundant roles in the cell autonomous response of microglia to FXN deficiency. Both NLRP2 and NLRP3 inflammasomes play major roles in response to damage-associated molecular patterns (DAMPs) associated with cell injury. FXN-deficient microglia may therefore be cell autonomously ‘primed’ for a multi-pronged inflammatory response to intrinsic cellular stress. Several top upregulated DEGs in FRDA microglia may generate cell intrinsic DAMPs via dysregulated ROS production, perturbations of intracellular organelle homeostasis, dysregulated ATP release, or activation of interferon pathways, all of which can prime both NLRP2- and NLRP3-mediated inflammasomes. We have observed that FXN-deficient microglia are enriched for the activated p33:p12 CASP1 dimers that remains bound to the inflammasome complex, with no detectable further p33 processing to generate the p20 subunit that may limit inflammasome activation (Boucher et al., 2018; Sandstrom and Vance, 2018). Exposure to non-cell intrinsic danger signals, including the presence of degenerating neurons, may sustain inflammasome activation but also lead to further processing of p33 to p20, thus modulating inflammasome responses.

Consistent with the activation of inflammasome-driven pathways, we detected increased secretion by FRDA microglia of both IL1B and IL6. Increased levels of these cytokines are hallmarks of inflammasome activation in response to viral infections, and in autoimmune diseases. Components of both the classical (C1R and C1S, both under-secreted) and alternative (CFD, oversecreted) complement pathways are also differentially represented in the secretome of FRDA vs. isogenic control microglia. Interesting, similar changes in complement components are observed in autoimmune diseases like systemic lupus erythematosus and rheumatoid arthritis (Jia et al., 2022; Ma et al., 2025; McMurray et al., 2024). Furthermore, other over- and under-secreted proteins by FRDA microglia are associated with autoimmune conditions, including CHI3L1 (Deng et al., 2025), uPAR/PLAUR (Dinesh and Rasool, 2018), LBP (Losanto et al., 2021), and MRC1 (Zhang et al., 2023), to name only a few. This finding suggests that the characterization of differentially secreted proteins in FXN-deficient microglia might identify candidate blood and cerebrospinal fluid biomarkers for FRDA, some of which may be shared with autoimmune diseases.

Although NLRP2 and NLRP3 may have overlapping functions in FRDA microglia, NLRP2 is also likely to have distinct roles independent of inflammasome activation. In oocytes and preimplantation embryos, NLRP2 functions as a maternal-effect protein within the subcortical maternal complex, where it regulates DNA methylation at imprinted loci and supports zygotic genome activation through protein-protein interactions that control the localization of DNA methyltransferases and other regulatory proteins (Amoushahi et al., 2019; Sharif et al., 2026). Consistent with this role, maternal deletion of *Nlrp2* in mice alters both the methylome and transcriptome of germinal vesicle oocytes, with impaired post-transcriptional regulation thought to underlie many of the transcriptional changes (Anvar et al., 2025). Our present study identified two other maternal effect genes among the top up-regulated DEGs in FRDA microglia, namely *MEG3* and *MEG8*, which also have roles in the regulation of DNA methylation and gene expression (Farhadova et al., 2024; Terashima et al., 2018). Together, these findings raise the possibility that coordinated dysregulation of NLRP2 and other epigenetic regulators contributes to transcriptomic remodeling and inflammatory pathways in FRDA through mechanisms distinct from those mediated by NLRP3.

In conclusion, our findings support the concept that microgliopathy represents an important contributor to neurodegeneration in FRDA. At early stages of disease progression, possibly even prior to the onset of neuronal degeneration, FXN-deficient microglia might respond to cell intrinsic danger signals by becoming overactive, which in turn might cause the reactivation of neighbouring astrocytes. Together, reactive astrocytes and activated microglia might establish a neuroinflammatory microenvironment that might begin to affect the survival of FXN-deficient neurons even before cell-intrinsic neuronal cell death pathways are activated. At later stages, when the degeneration of FXN-deficient neurons becomes more prominent, microglia might be confronted with new, neuron-derived, danger signals that further modify their pathobiological state, possibly causing further activation of neuroinflammatory mechanisms aimed at removing dying neurons. These possible scenarios suggest that targeting activated microglia early in the progression of FRDA might offer a previously unrecognized early window of intervention. The dysregulated genes and proteins identified in this study might offer attractive targets for such therapeutic strategies.

## Materials and Methods

### Human induced pluripotent stem cells

The following human iPSC lines were used: FRDA_86 (female donor) and FRDA_86 GAA homo-del, FRDA_4676 (male donor) and FRDA_4676 GAA homo-del, FRDA_4259 (male donor) and FRDA_4259 GAA homo-del. All lines were obtained from the Friedreich’s Ataxia Cell Line Repository (https://labs.utsouthwestern.edu/napierala-lab/cell-line-repository).

Undifferentiated state of human iPSCs was assessed by testing for expression of the stem cell markers NANOG and OCT4 using rabbit anti-NANOG (1/1,000; Abcam; Cambridge, UK; Cat. No. ab21624) and rabbit anti-OCT4 (1 μg/ml; Abcam; Cat. No. ab19857) or goat anti-OCT3/4 (1/500; Santa Cruz Biotechnology; Dallas, TX, USA, Cat. No. sc-8628) antibodies, as previously described (Thiry et al., 2025).

### Microglia derivation from human iPSCs

Derivation of microglia from human iPSCs was performed following two previously published protocols (Haenseler et al., 2017, Douvaras et al., 2024). All iPSC-derived microglia preparations were validated through morphological analysis, immunocytochemistry, and RNAseq as described (Tang et al., 2022; Thiry et al., 2025).

### RNA sequencing

For bulk RNAseq, total RNA was isolated from cell pellets obtained from iPSC-derived microglia using sequential treatment with TRIzol Reagent (ThermoFisher Scientific; Cat. No. 15596026) and PureLink RNA Micro Scale Kit (ThermoFisher Scientific; Cat. No. 12183-016) following the instructions provided by the manufacturer. RNA samples were isolated from eight iPSC-derived microglia preparations (two female and two male isogenic microglia samples, two female and two male FRDA microglia samples). All RNA samples were analyzed by Illumina next-generation sequencing at the Genomics platform at the Institute for Research in Immunology and Cancer, Montreal, Quebec, Canada (https://www.iric.ca/en/research/platforms-andinfrastructures/genomics). Adaptor sequences and low-quality bases in the resulting FASTQ files were trimmed using Trimmomatic version 0·35 (Bolger et al., 2014), and genome alignments were conducted using STAR version 2·5 1b (Dobin et al., 2013). Sequences were aligned to the human genome version GRCh38 using STAR version 2.7.1a, with gene annotations from Gencode v40 based on Ensemble release 106. As part of quality control, the sequences were aligned to several different genomes to verify that there was no sample contamination. Raw read-counts were obtained directly from STAR and reads in transcripts per million (TPM) formats were computed using RSEM (Li and Dewey, 2011).

### In silico analysis

To characterize iPSC-derived microglia preparations by RNAseq, the levels of expression of 49 *bona fide* microglia/macrophage marker genes (Butovsky et al., 2014; Hickman et al., 2013) were compared in induced microglia *versus* undifferentiated iPSCs (open-source data). Read counts for these specific genes were converted into log2-counts-per-million (logCPM) values as previously described (Tang et al., 2022). Accordingly, negative values represent very low gene expression values. Differential gene expression analysis was conducted on the raw read-count matrix using DESeq2 version 1.30.1 (Love et al., 2014). The experimental batch and sex were incorporated into the design used for differential expression analysis since they were the two known sources of variability in the studied samples. DEGs were considered statistically significant if the adjusted p-value (false discovery rate; FDR) was <0.05, with no fold-change cut-off applied. Over-representation Analysis (ORA) was used to functionally characterize DEGs. ORA identifies biological processes or pathways that are statistically over-represented among a predefined list of significant DEGs (based on adjusted p-value and fold-change thresholds). This approach evaluates whether certain functional gene sets occur more frequently than expected by chance within the subset of significantly up- or down-regulated genes. GO terms and KEGG pathways over-represented in the DEGs were computed separately for all up- and all down-regulated genes. All genes expressed in the dataset were used as the background universe and terms with a Bonferroni-correct p value < 0.05 were reported.

### Open-source data

Raw read counts from RNA sequencing of undifferentiated human iPSCs were obtained from the LINCS Data Portal (LSC-1002 and LSC-1004 from Dataset LDS-1355) (http://lincsportal.ccs.miami.edu/datasets/). LSC-1002 and LSC-1004 correspond to human iPSC lines CS14i-CTR-n6 (female donor) and CS25iCTR-18n2 (male donor), provided by Cedar Sinai Stem Cell Core Laboratory. These raw data were combined with the RNAseq data from the present microglia preparations and analyzed using a single pipeline, to generate heatmaps comparing the levels of expression of activated microglia marker genes in each sample.

### Secretome analysis of microglia conditioned media

Conditioned media from multiple iPSC-derived microglia preparations generated at different timepoints were subjected to analysis of 285 secreted proteins utilising the nucleobase Enabled Localized Immunoassay with Spectral Addressing (nELISA) platform from Nomic-Bio (Montreal, Quebec, Canada; https://www.nomic.bio/) (Dagher et al., 2025). Briefly, antibody pairs were pre-assembled on spectrally encoded microparticles (Dagher et al., 2018), resulting in spatial separation between non-cognate antibodies, preventing the rise of reagent-driven cross-reactivity, and enabling multiplexing of hundreds of ELISAs in parallel using flow cytometry. Standard curves were included for each protein, enabling quantification of pg/mL concentrations.

### Protein expression studies

Western blotting was performed using the following antibodies: rabbit anti-NLRP2 (Proteintech; 15182-1-AP), mouse anti-NLRP3 (Adipogen; AG-20B-0014-C100), rabbit anti-ASC (Adipogen; AG-25B-0006-C), rabbit anti-CASP1 (Abcam; ab207802, rabbit anti-CASP1 p10/p12 (Invitrogen; MA5-56516), rabbit anti-LAMP1 (Abcam; ab24170), mouse anti-Rab7 (Abcam; ab50533), and mouse anti-FXN (Abcam; 110328). Cross-linking using DTSSP reagent was performed as described (Palaparti et al., 1997), followed by non-reducing SDS-PAGE. All protein expression studies were performed with at least 3 biologically independent preparations per condition. Error bars shown in figures are means ± standard error of means (SEM) of the average. Unpaired t test was used to detect significant differences in protein expression levels.

## Data Availability Statement

The raw data are available at the Sequence Read Archive (SRA) (accession number: PRJNA1472680). Data are accessible using the following link:

https://can01.safelinks.protection.outlook.com/?url=https%3A%2F%2Fwww.ncbi.nlm.nih.gov%2Fsra%2FPRJNA1472680&data=05%7C02%7Clouise.thiry%40mcgill.ca%7C58d83b81a0264ac298b608debe446120%7Ccd31967152e74a68afa9fcf8f89f09ea%7C0%7C0%7C639157398079968335%7CUnknown%7CTWFpbGZsb3d8eyJFbXB0eU1hcGkiOnRydWUsIlYiOiIwLjAuMDAwMCIsIlAiOiJXaW4zMiIsIkFOIjoiTWFpbCIsIldUIjoyfQ%3D%3D%7C0%7C%7C%7C&sdata=TPPTHaePEXSg5pH6MJLTuEPbMnmIX%2BZKPfHzJJQ1xIw%3D&reserved=0

## Conflict of Interest Statement

The authors declare that they have no conflict of interest.

## Author Contributions

YT, RL performed experiments. LT, MF performed data analysis. LT prepared figures and wrote part of the manuscript. SS and SF supervised lab members. SS and MP conceived the study plan. SS supervised the studies, data analysis, and manuscript preparation.

## Acknowledgments

We thank Vincent Soubannier, Valerio Piscopo, and Christian Rampal for help at various stages. We also thank Dr. Peter McPherson for providing anti-Rab7 antibody. These studies were funded by the Friedreich’s Ataxia Research Alliance and the Christina Foundation. SS is a Distinguished James McGill Professor of McGill University.

## Supplemental Information

List of all statistically significant DEGs

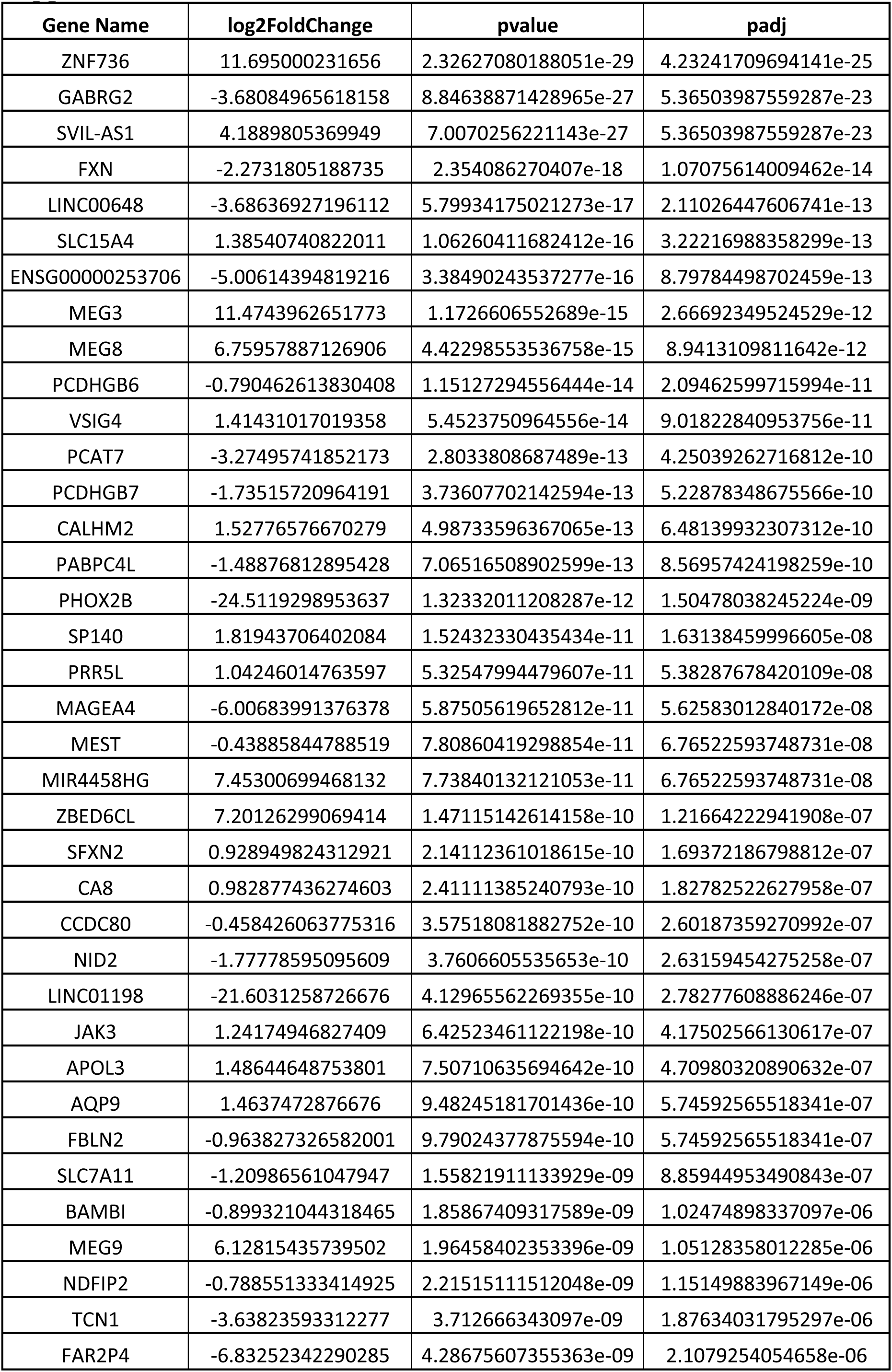
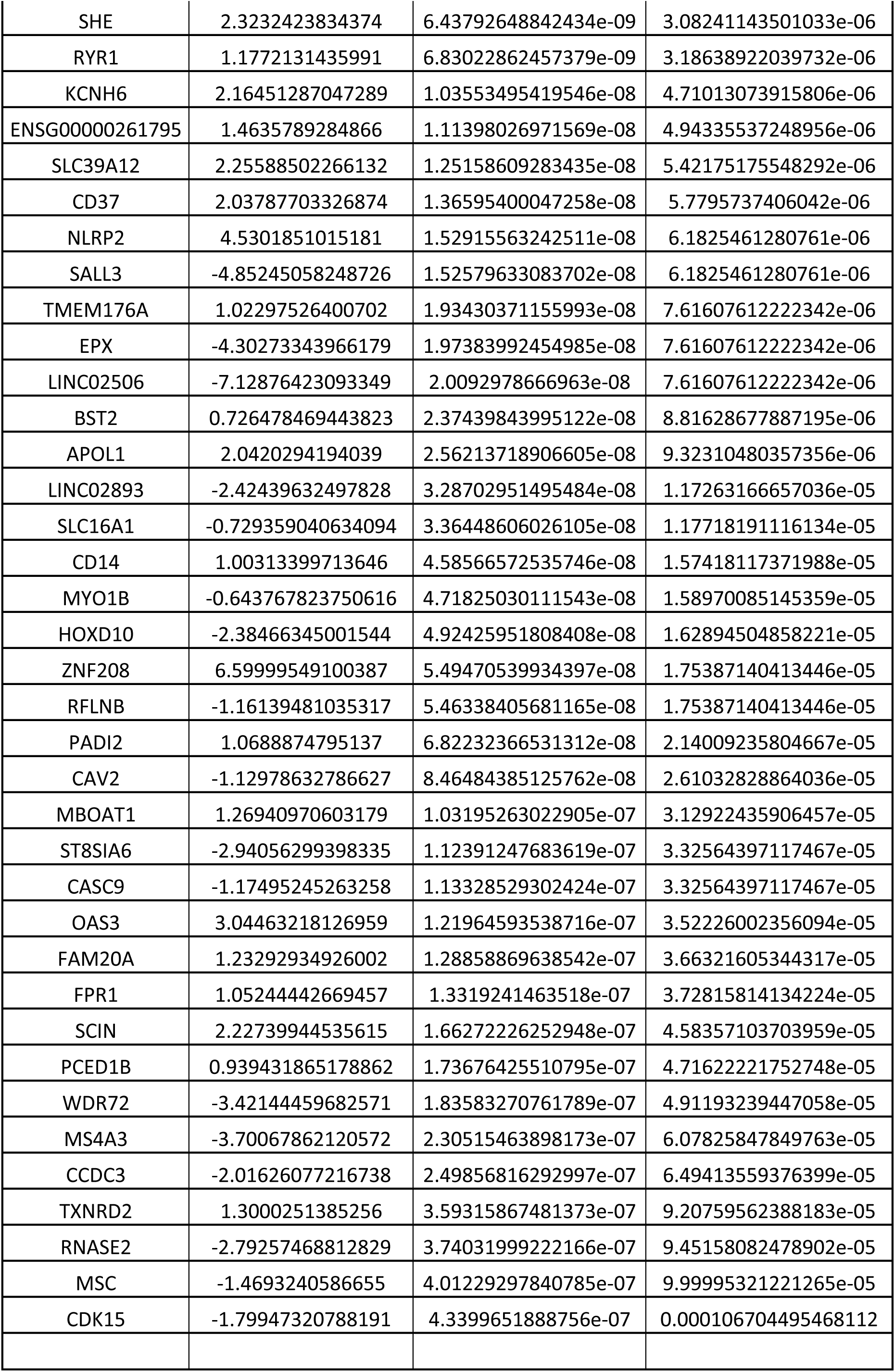
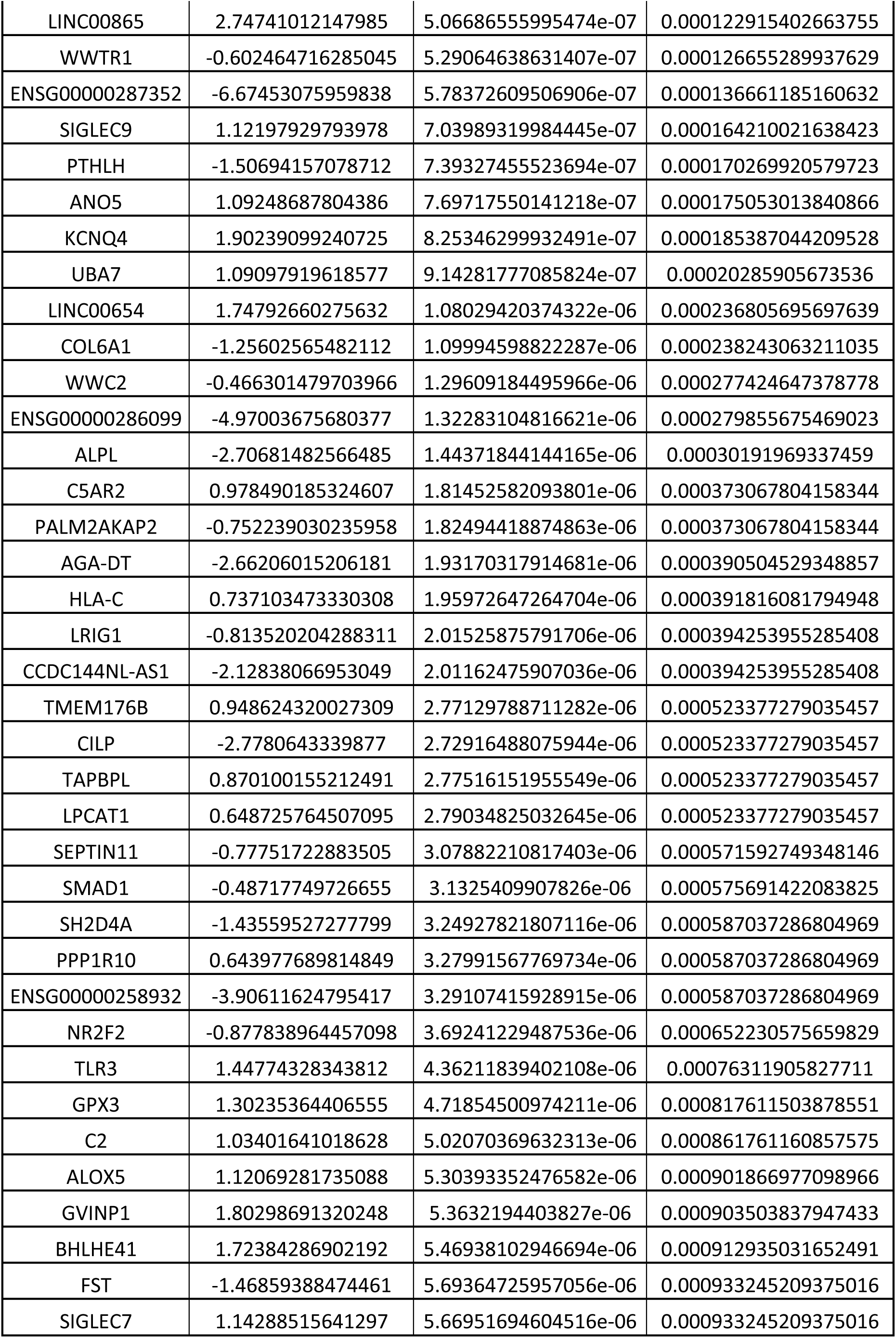
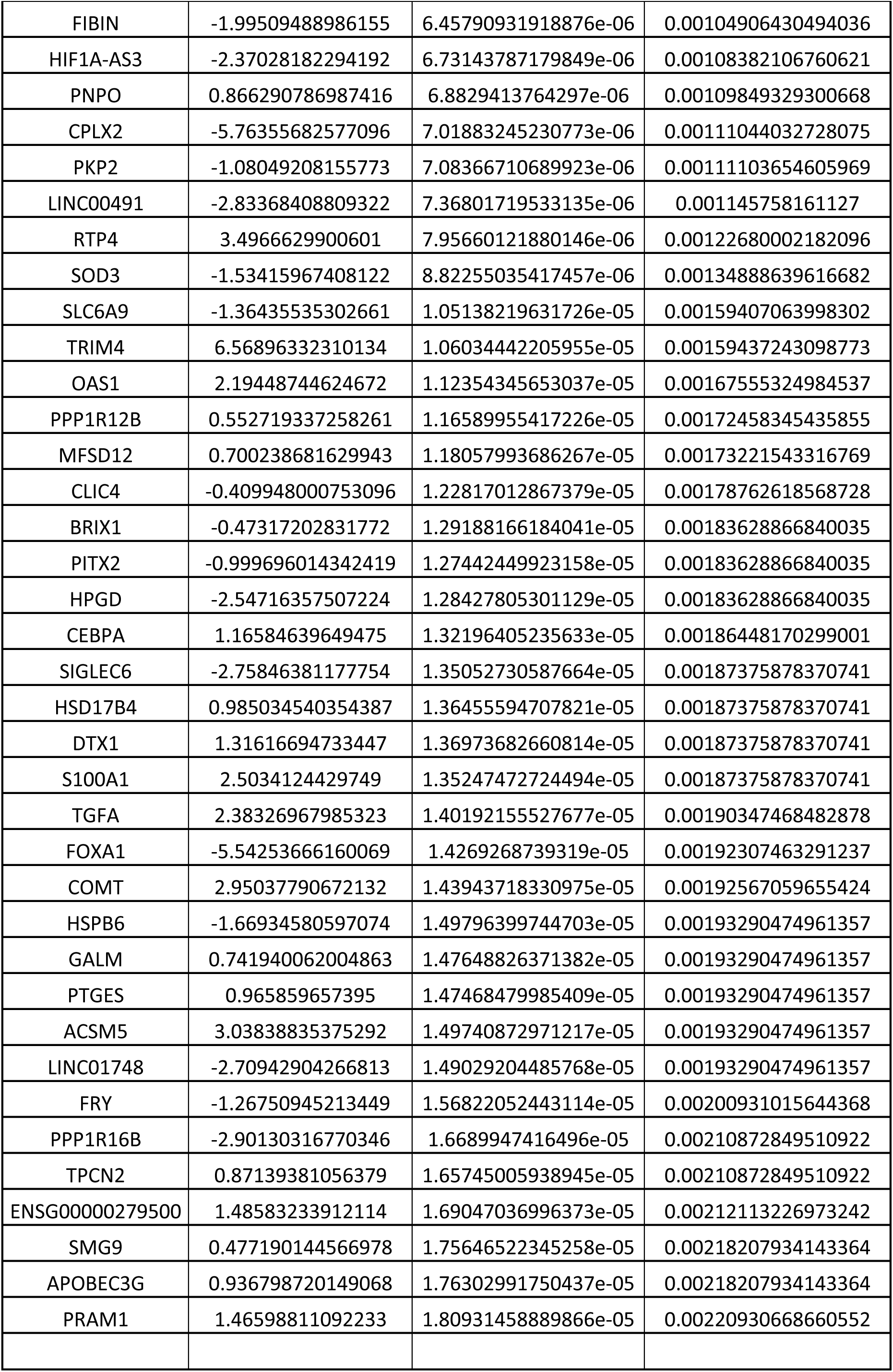
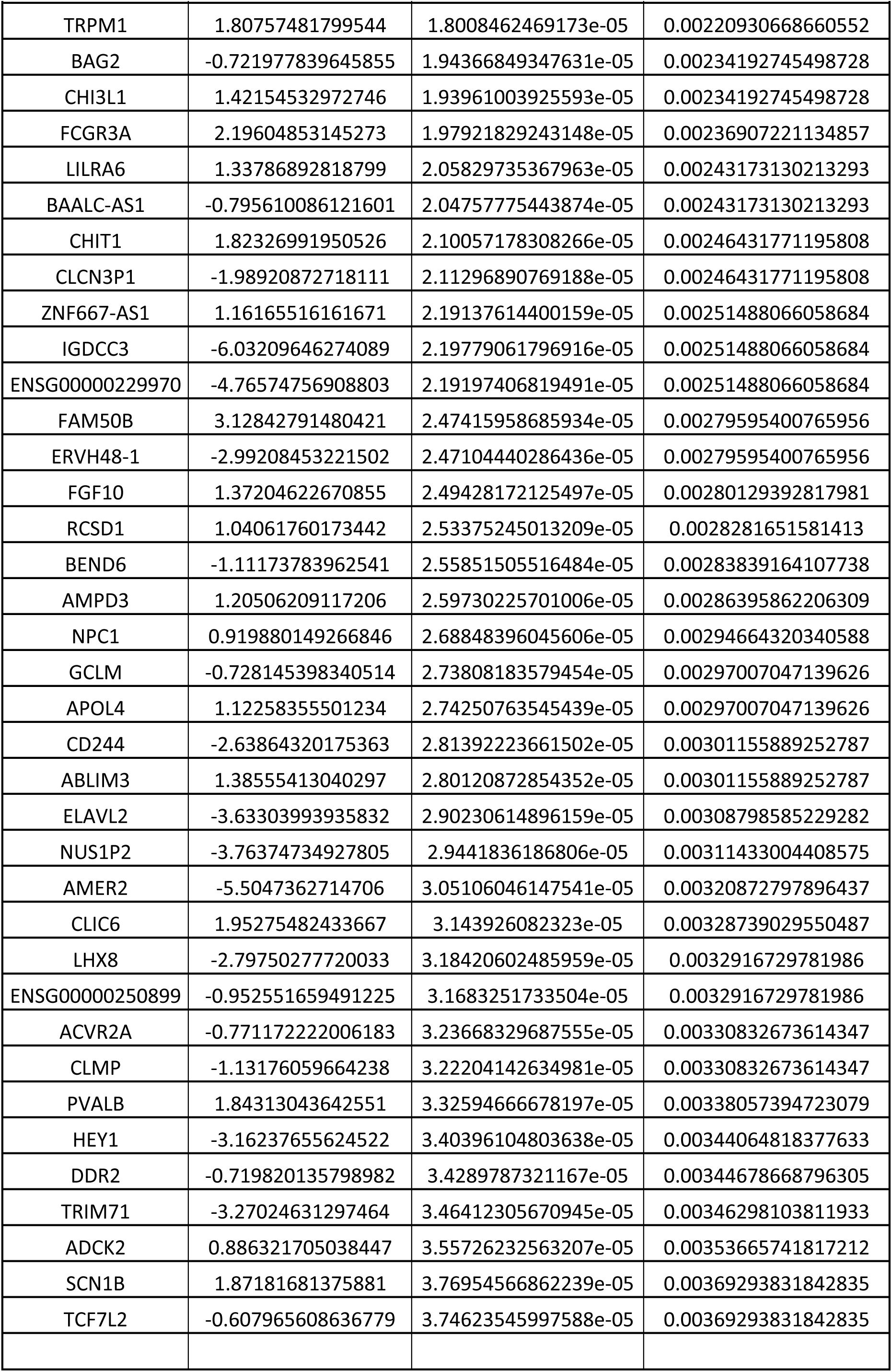
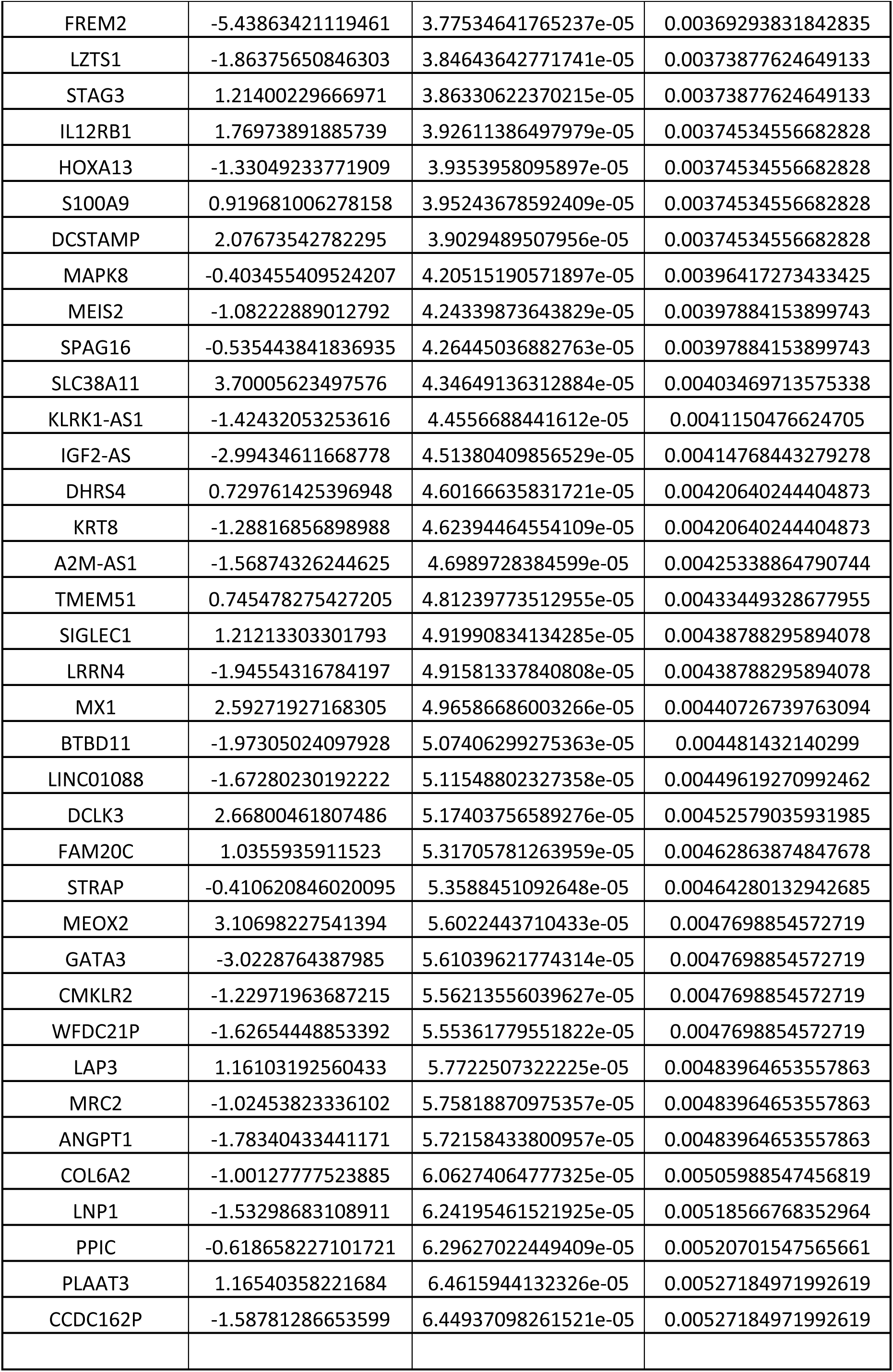
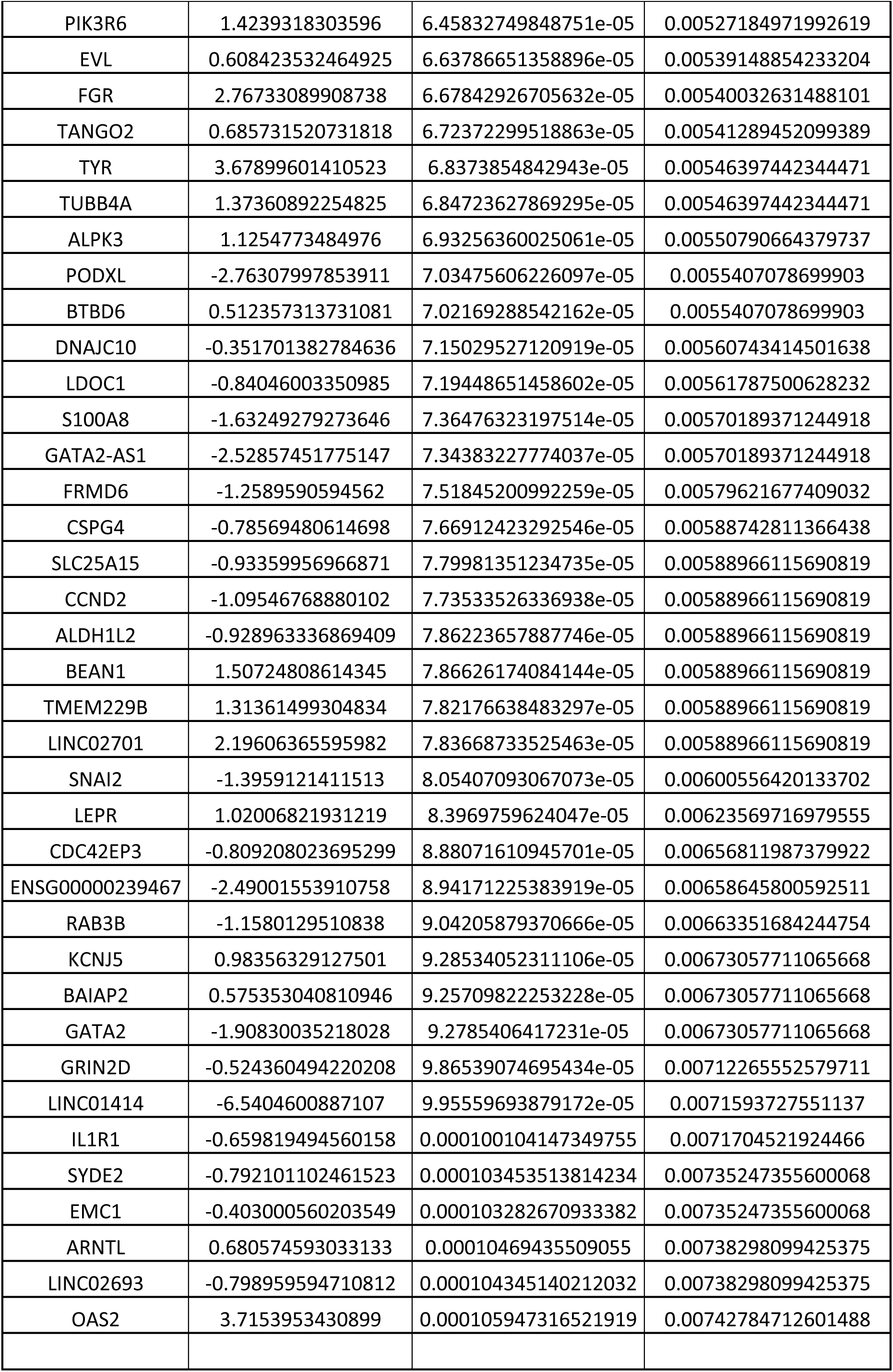
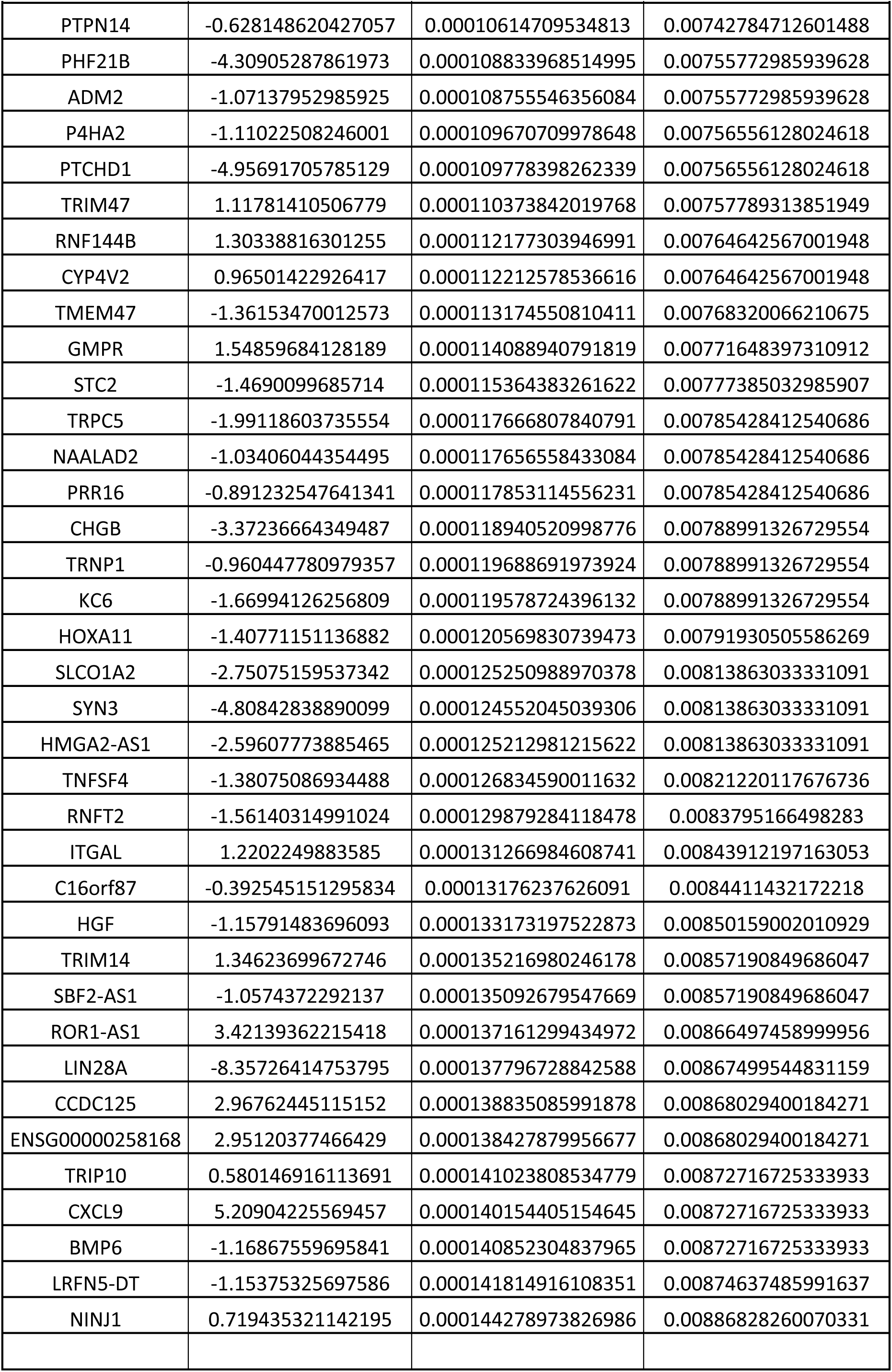
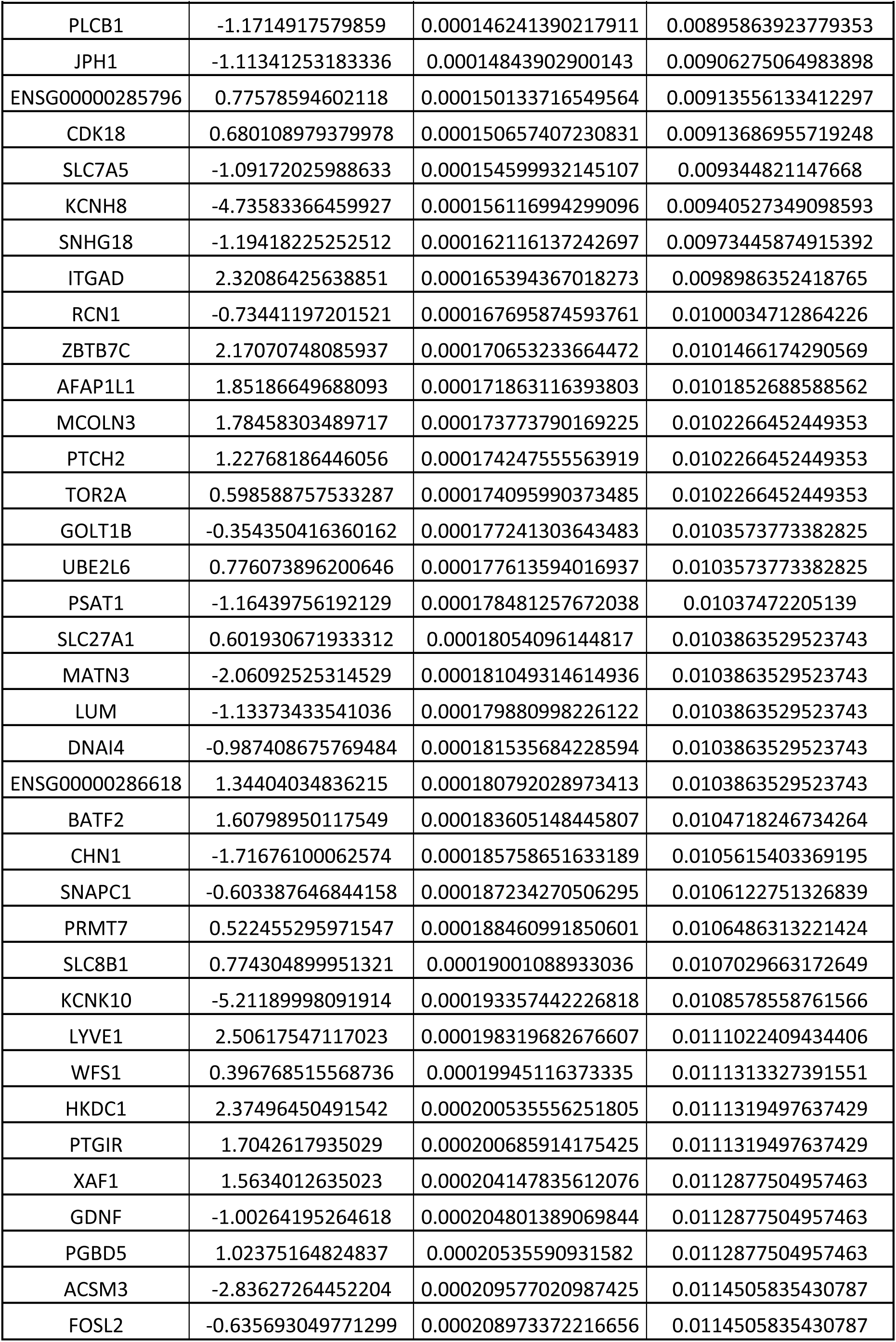
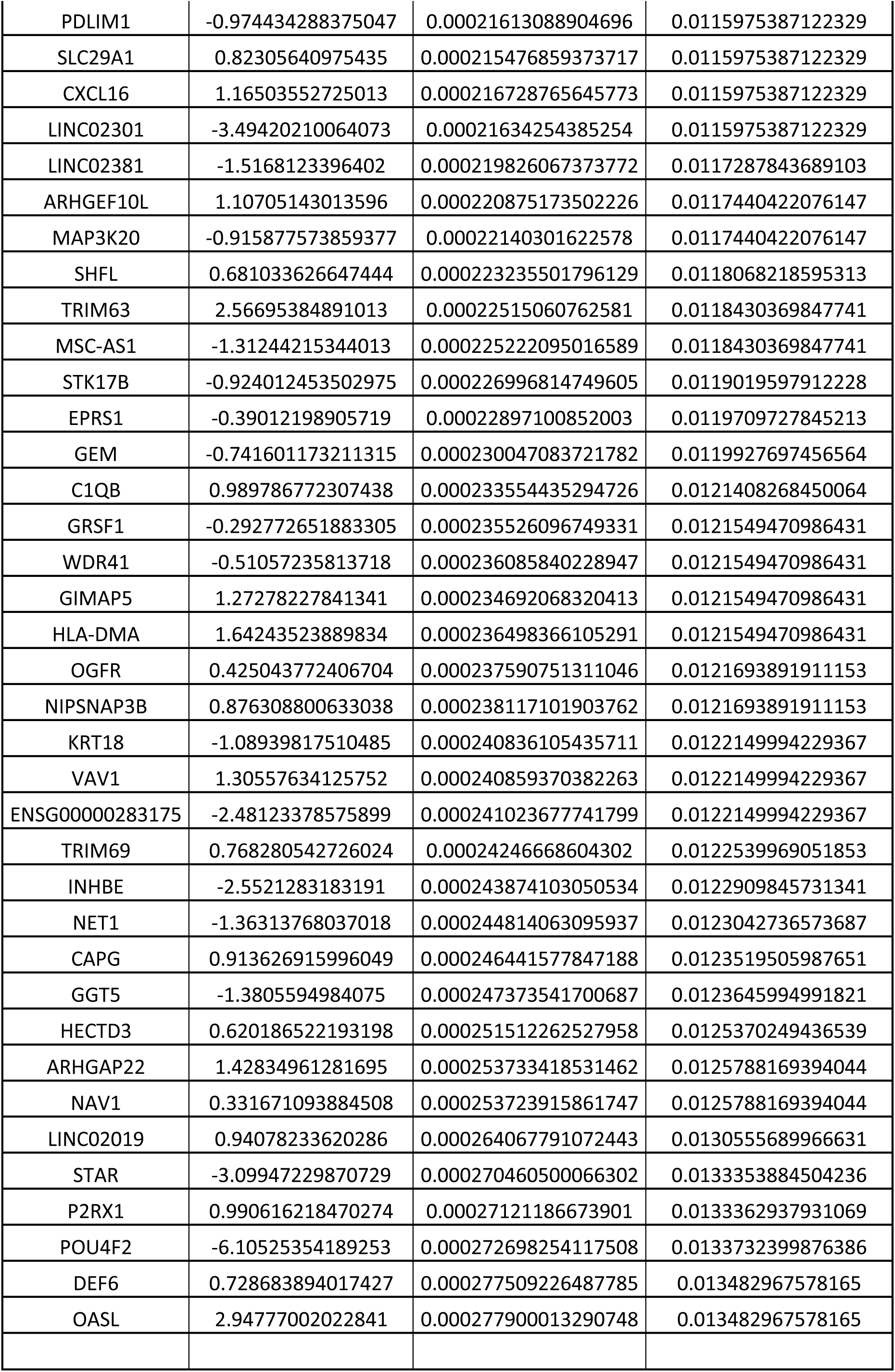
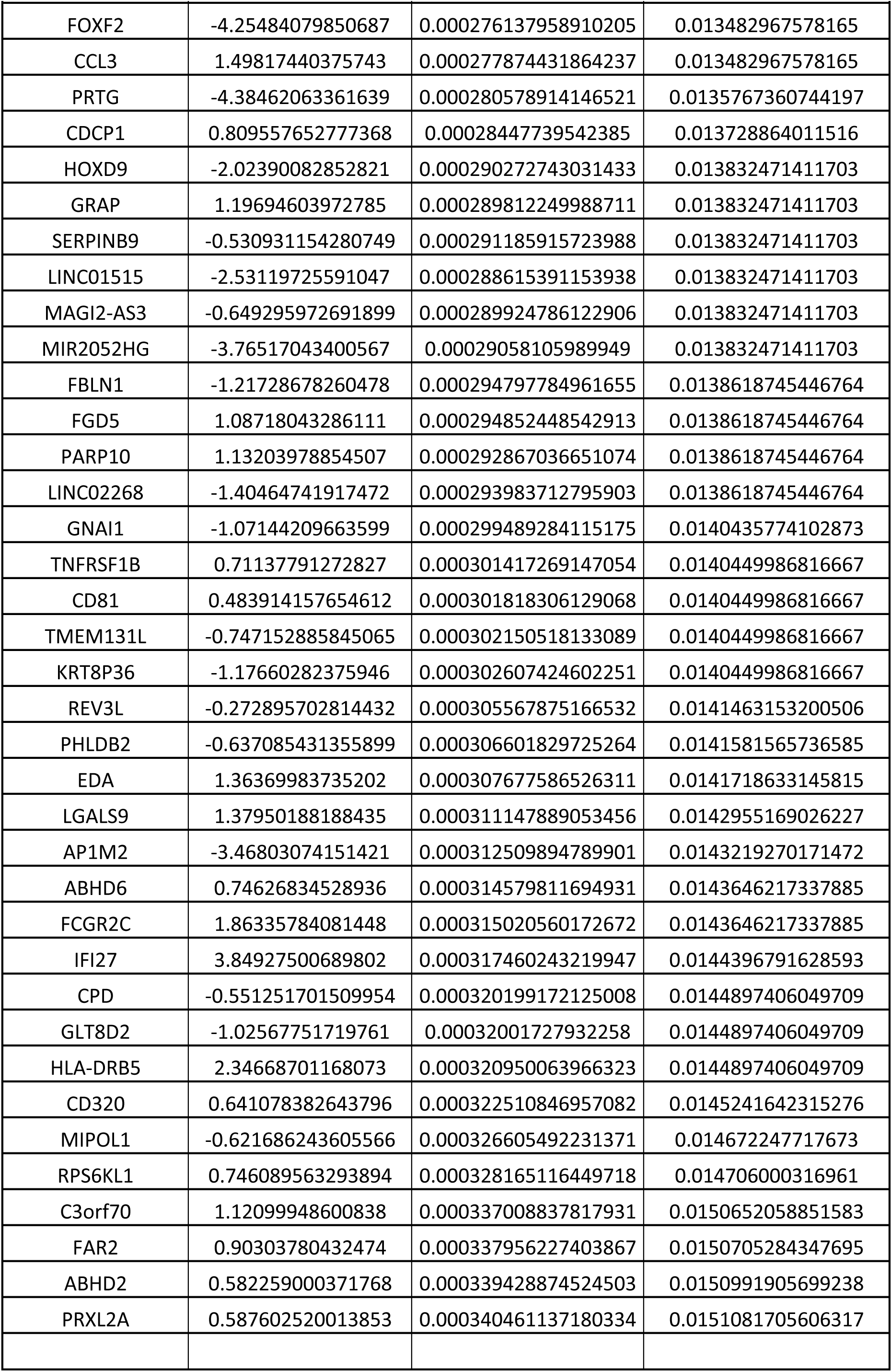
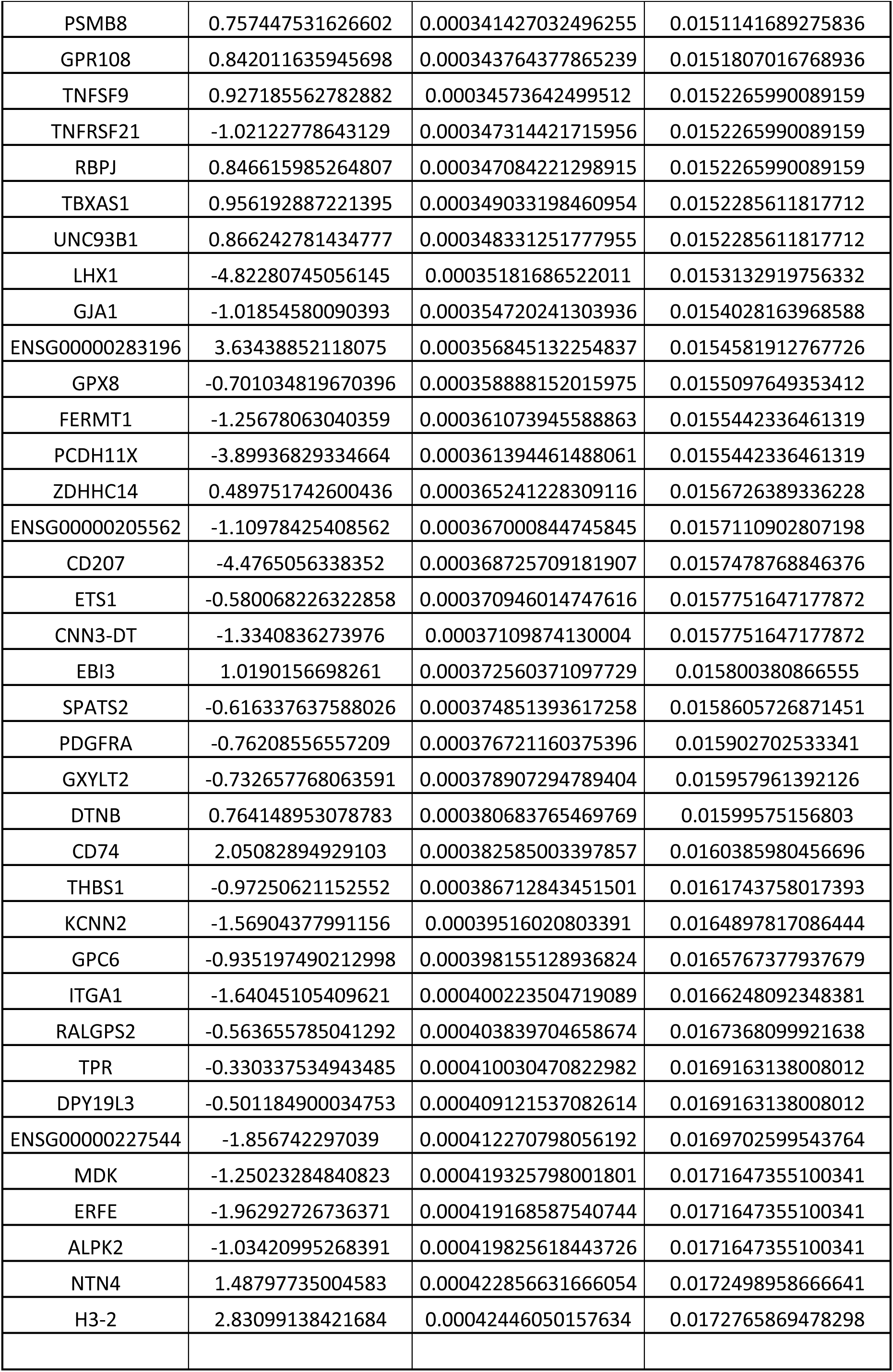
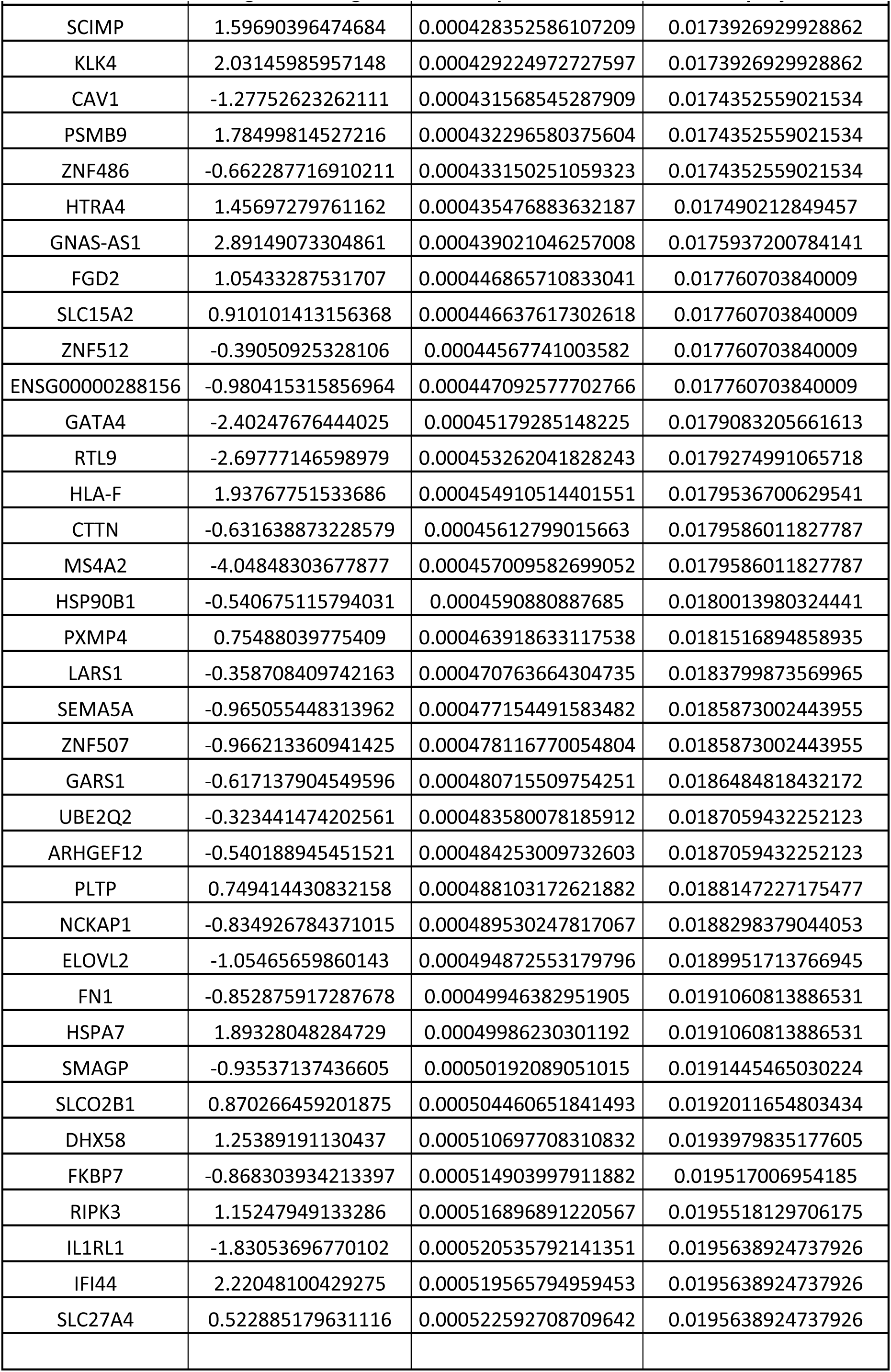
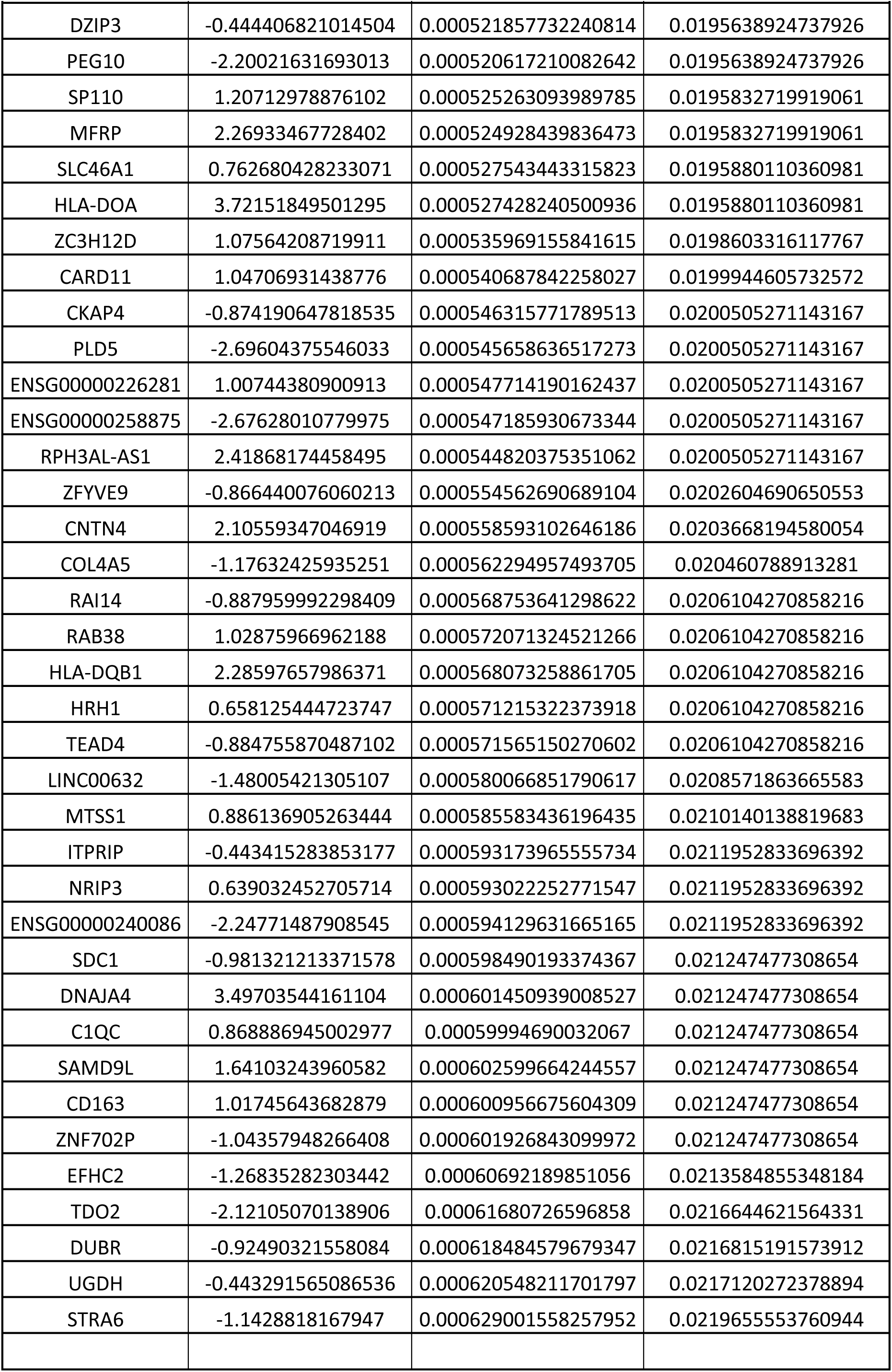
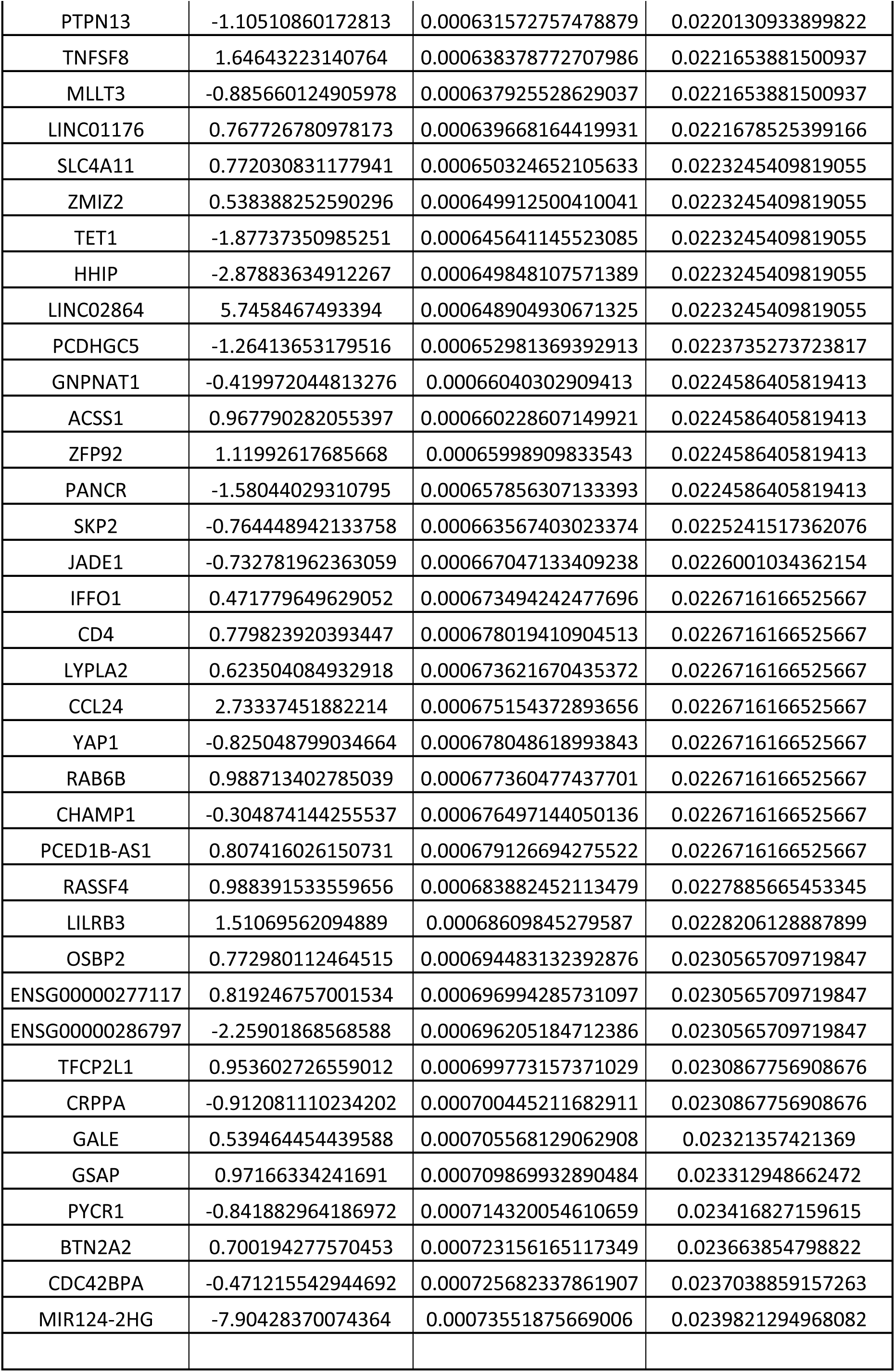
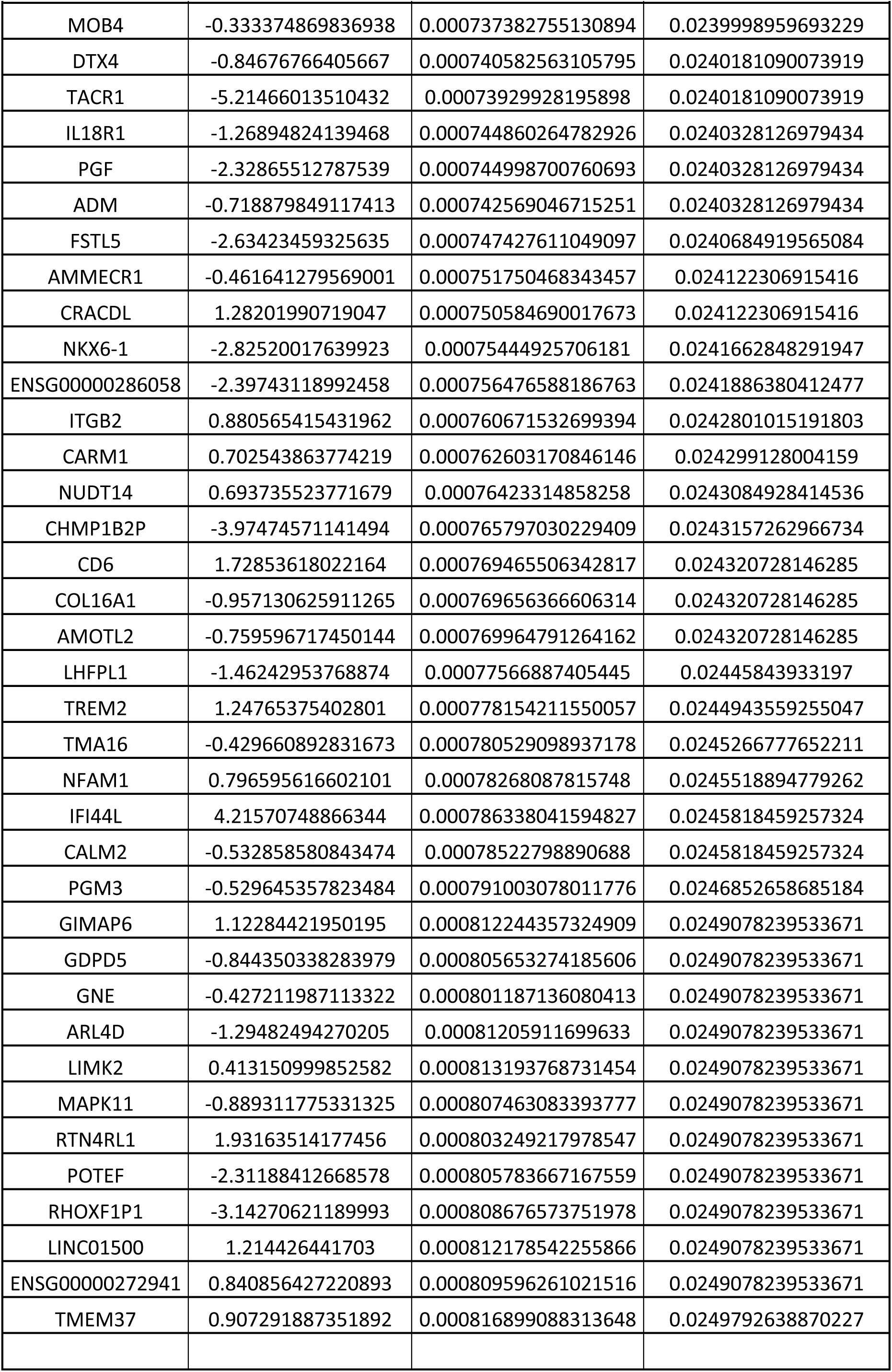
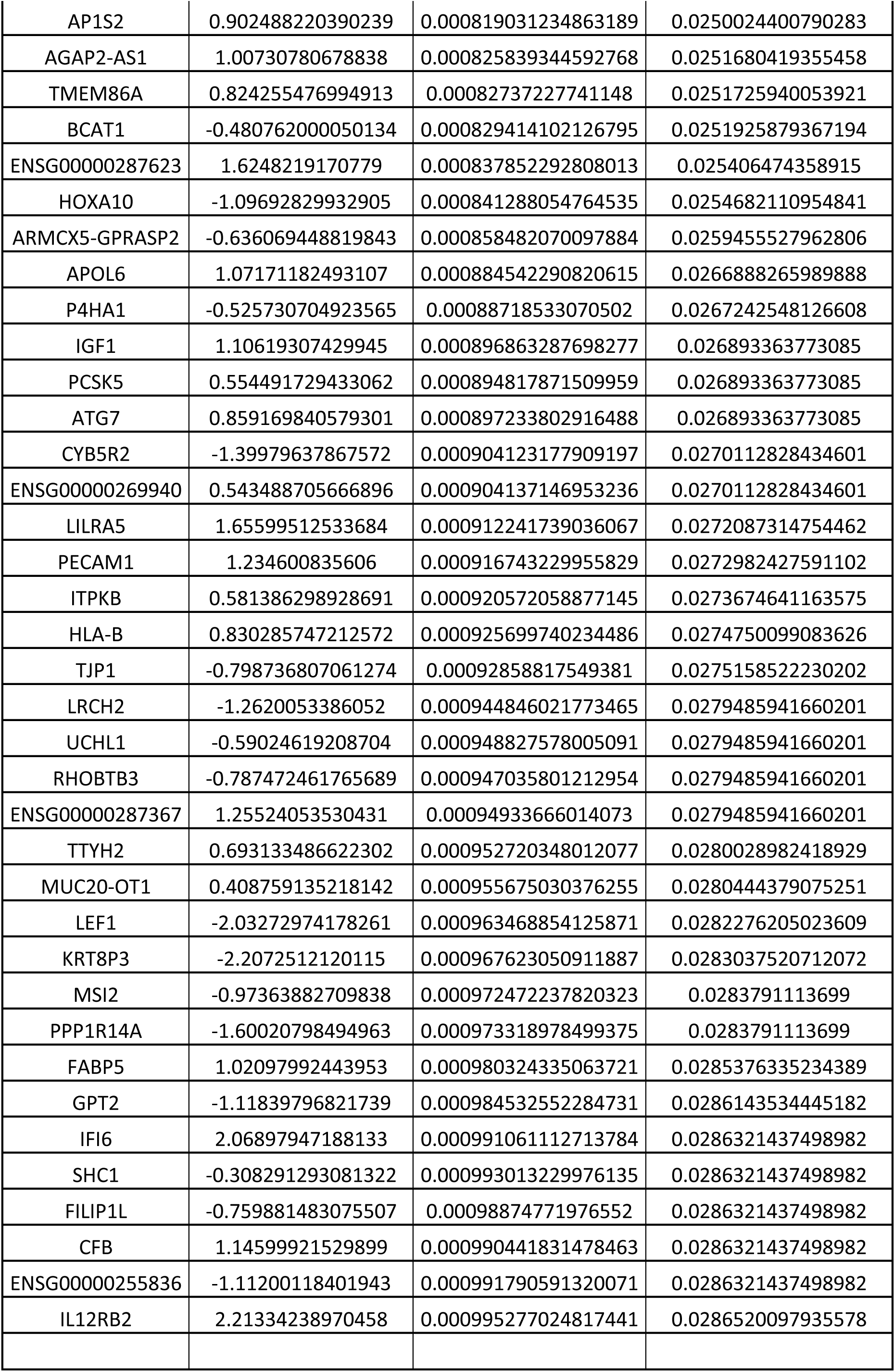
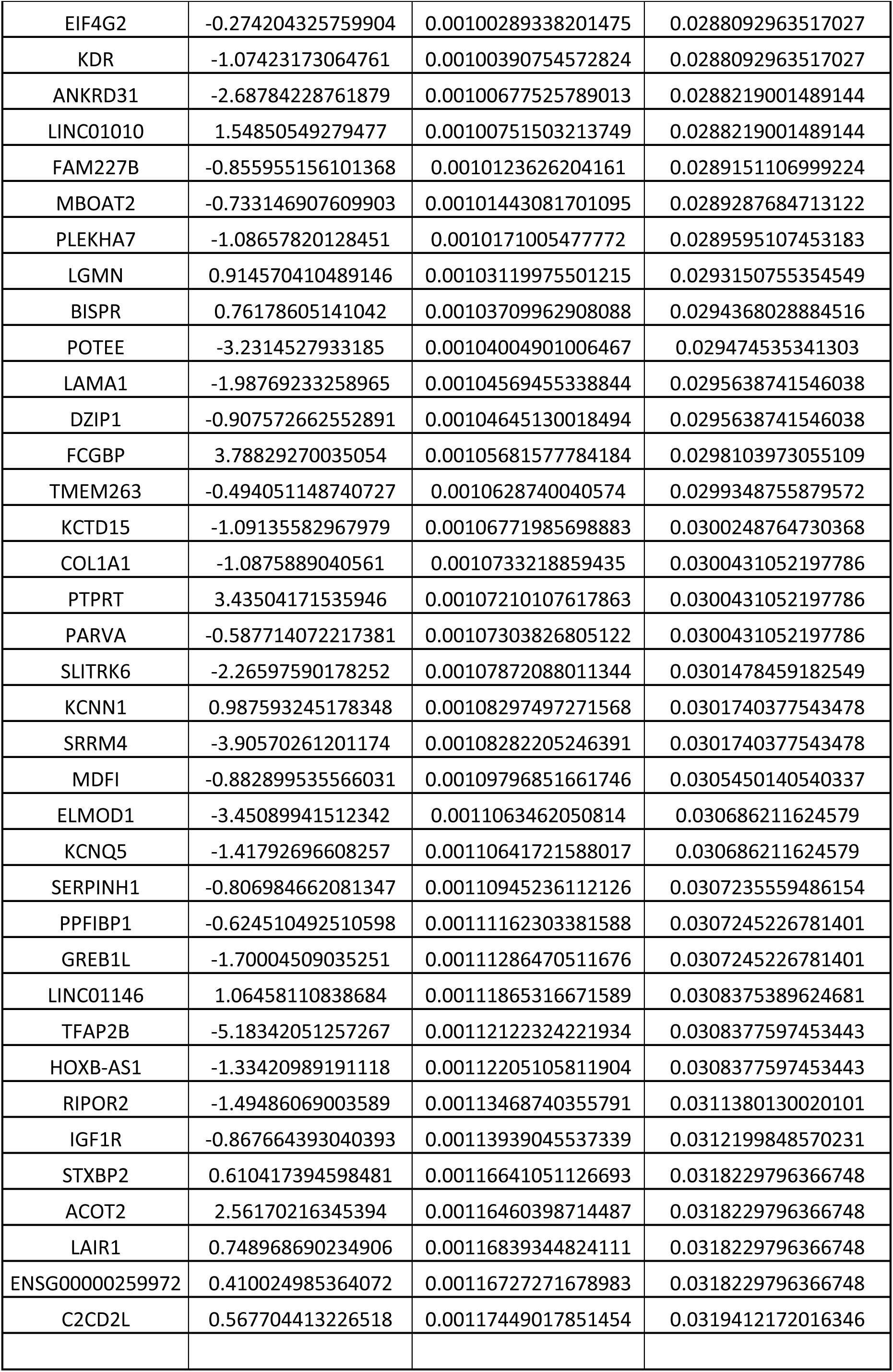
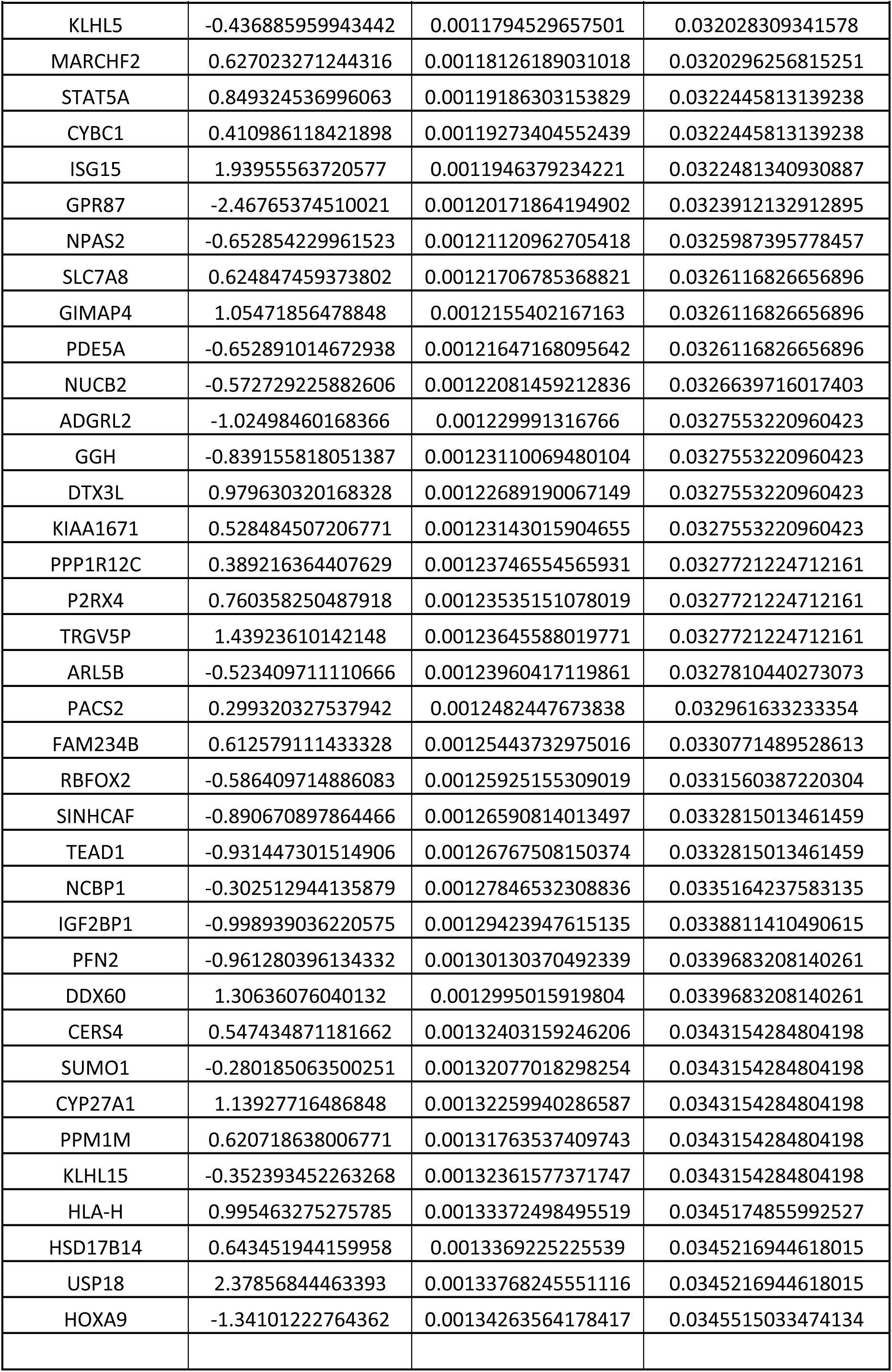
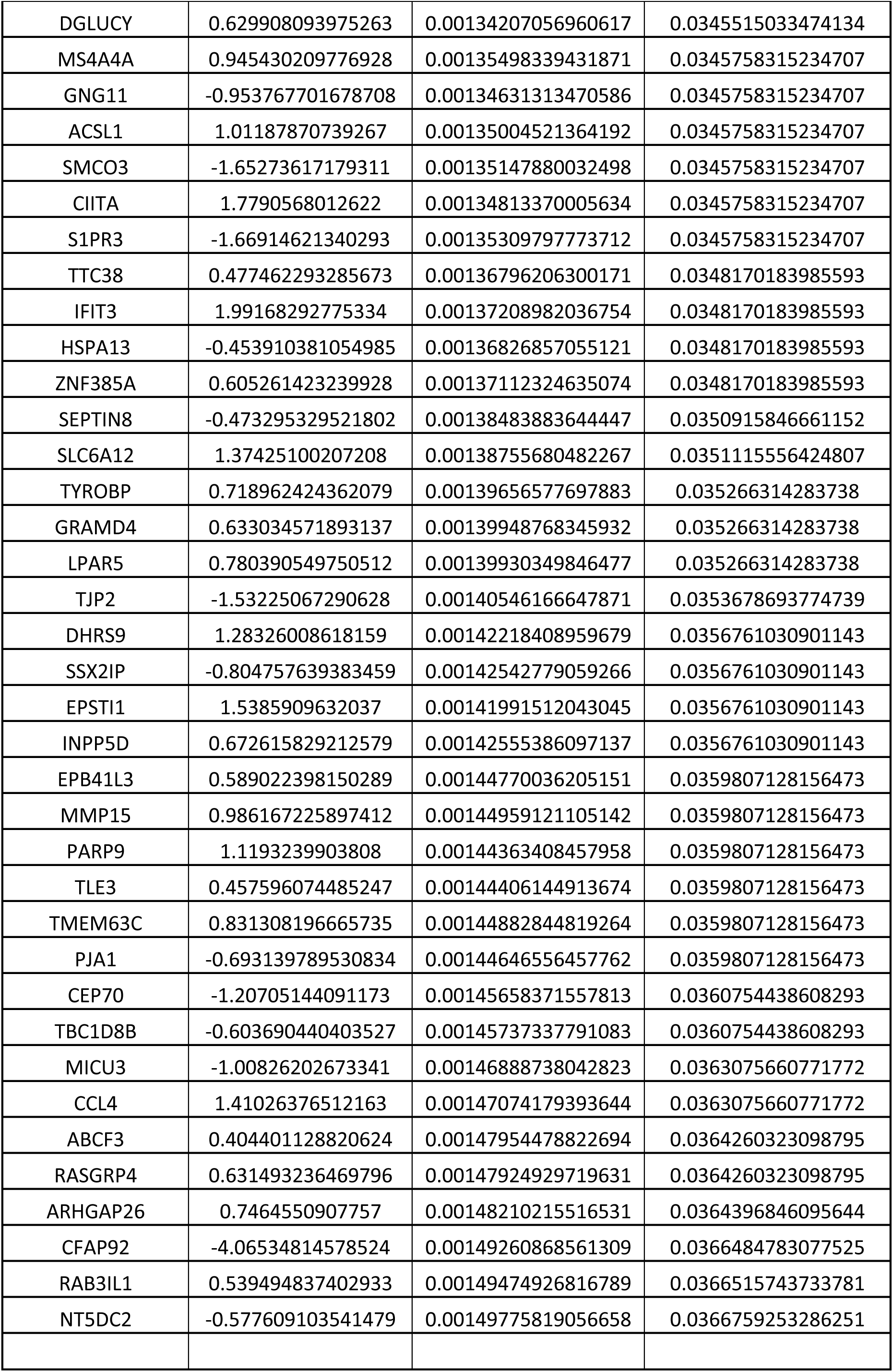
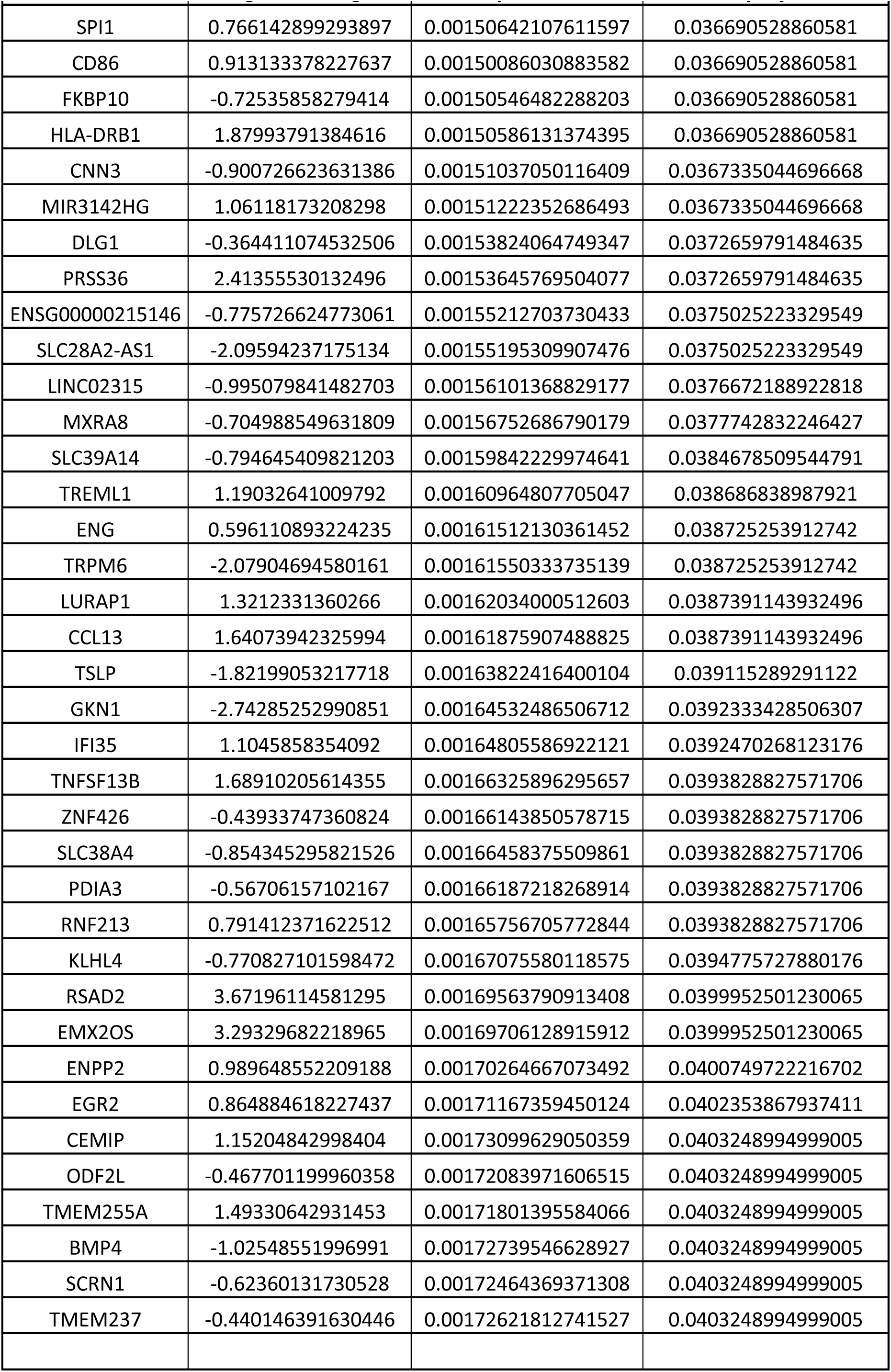
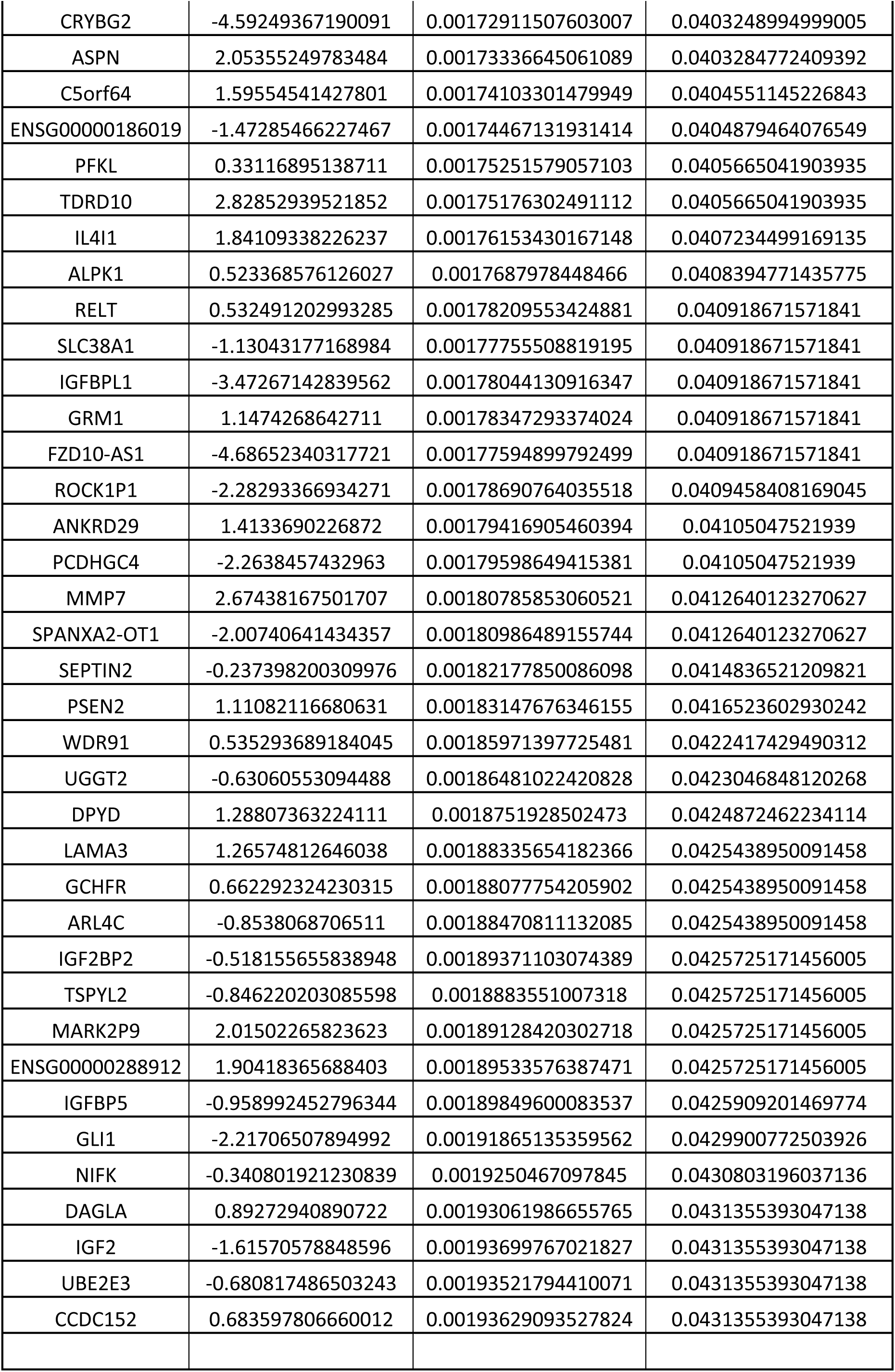
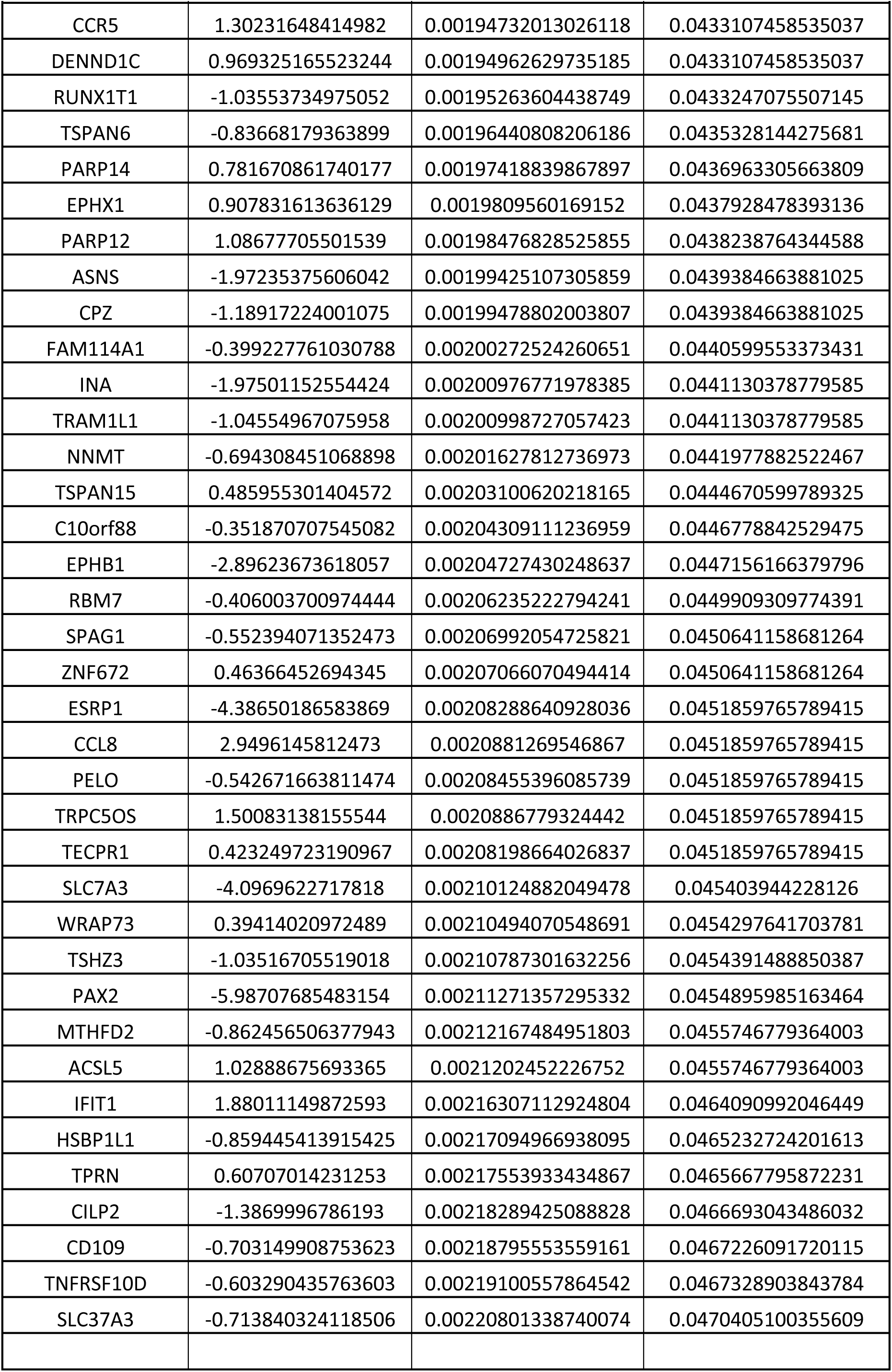
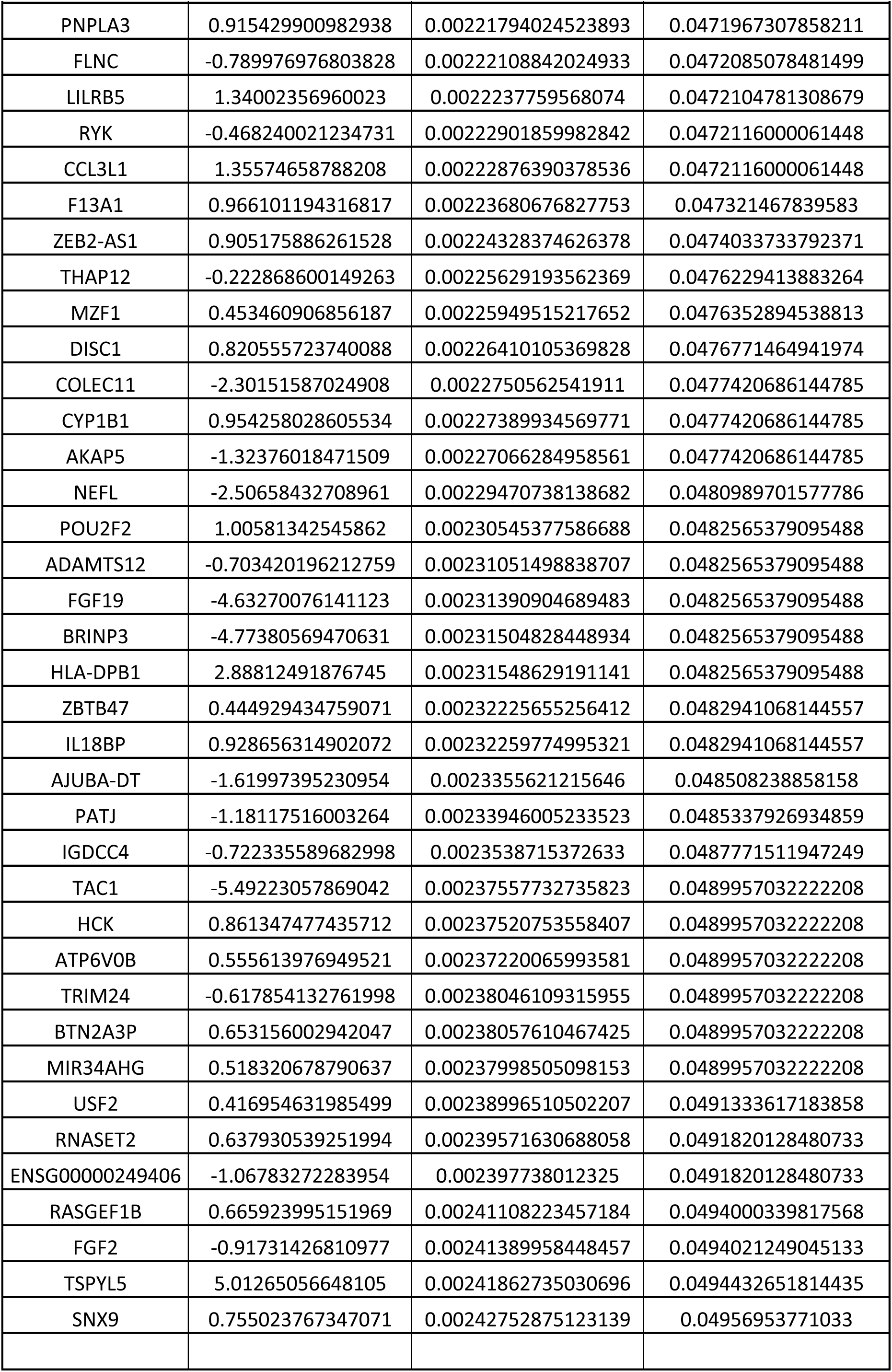
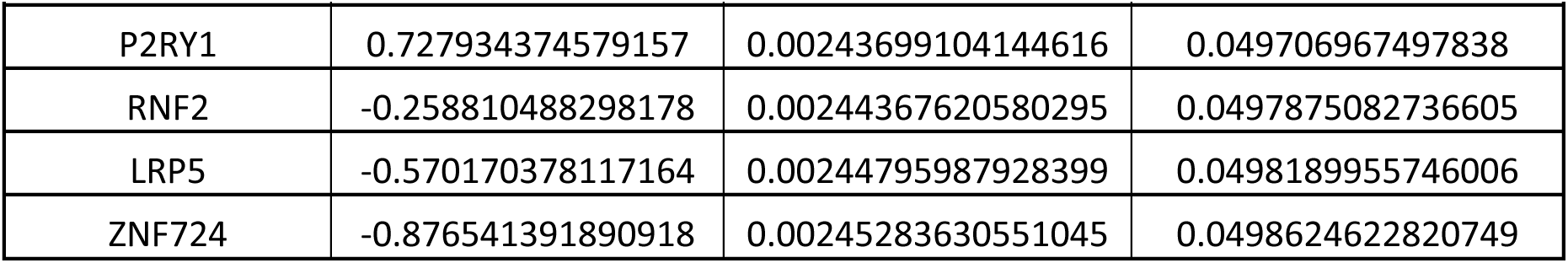

## Notes

### Competing Interest Statement

The authors have declared no competing interest.

## References

Albalawi F, Lu W, Beckel JM, Lim JC, McCaughey SA, Mitchell CH. The P2X7 Receptor Primes IL-1β and the NLRP3 Inflammasome in Astrocytes Exposed to Mechanical Strain. Front Cell Neurosci. 2017 Aug 8;11:227.

Amoushahi M, Sunde L, Lykke-Hartmann K. The pivotal roles of the NOD-like receptors with a PYD domain, NLRPs, in oocytes and early embryo development†. Biol Reprod. 2019 Aug 1;101(2):284–296.

Anvar Z, Jochum MD, Chakchouk I, Sharif M, Demond H, To AK, Kraushaar DC, Wan YW, Mari MC, Andrews S, Kelsey G, Van den Veyver IB. Maternal loss of mouse Nlrp2 alters the transcriptome and DNA methylome in GV oocytes and impairs zygotic genome activation in embryos. Clin Epigenetics. 2025 Jun 3;17(1):92.

Becher B, Derfuss T, Liblau R. Targeting cytokine networks in neuroinflammatory diseases. Nat Rev Drug Discov. 2024 Nov;23(11):862–879.

Biasizzo M, Kopitar-Jerala N. Interplay Between NLRP3 Inflammasome and Autophagy. Front Immunol. 2020 Oct 9;11:591803.

Blazer A, Qian Y, Schlegel MP, Algasas H, Buyon JP, Cadwell K, Cammer M, Heffron SP, Liang FX, Mehta-Lee S, Niewold T, Rasmussen SE, Clancy RM. *APOL1* variant-expressing endothelial cells exhibit autophagic dysfunction and mitochondrial stress. Front Genet. 2022 Sep 27;13:769936.

Bolger AM, Lohse M, Usadel B. Trimmomatic: a flexible trimmer for Illumina sequence data. Bioinformatics. 2014 Aug 1;30(15):2114–20.

Boucher D, Monteleone M, Coll RC, Chen KW, Ross CM, Teo JL, Gomez GA, Holley CL, Bierschenk D, Stacey KJ, Yap AS, Bezbradica JS, Schroder K. Caspase-1 self-cleavage is an intrinsic mechanism to terminate inflammasome activity. J Exp Med. 2018 Mar 5;215(3):827–840.

Bufi AA, Di Stefano J, Papait A, Silini AR, Parolini O, Ponsaerts P. The central role of CXCL10-CXCR3 signaling in neuroinflammation and neuropathology. Cytokine Growth Factor Rev. 2025 Aug;84:20–34.

Butovsky O, Jedrychowski MP, Moore CS, Cialic R, Lanser AJ, Gabriely G, Koeglsperger T, Dake B, Wu PM, Doykan CE, Fanek Z, Liu L, Chen Z, Rothstein JD, Ransohoff RM, Gygi SP, Antel JP, Weiner HL. Identification of a unique TGF-β-dependent molecular and functional signature in microglia. Nat Neurosci. 2014 Jan;17(1):131–43.

Campuzano V, Montermini L, Moltò MD, Pianese L, Cossée M, Cavalcanti F, Monros E, Rodius F, Duclos F, Monticelli A, Zara F, Cañizares J, Koutnikova H, Bidichandani SI, Gellera C, Brice A, Trouillas P, De Michele G, Filla A, De Frutos R, Palau F, Patel PI, Di Donato S, Mandel JL, Cocozza S, Koenig M, Pandolfo M. Friedreich’s ataxia: autosomal recessive disease caused by an intronic GAA triplet repeat expansion. Science. 1996 Mar 8;271(5254):1423–7.

Campuzano V, Montermini L, Lutz Y, Cova L, Hindelang C, Jiralerspong S, Trottier Y, Kish SJ, Faucheux B, Trouillas P, Authier FJ, Dürr A, Mandel JL, Vescovi A, Pandolfo M, Koenig M. Frataxin is reduced in Friedreich ataxia patients and is associated with mitochondrial membranes. Hum Mol Genet. 1997 Oct;6(11):1771–80.

Chen Y, Qian J, Ding P, Wang W, Li X, Tang X, Tang C, Yang Y, Gu C. Elevated SFXN2 limits mitochondrial autophagy and increases iron-mediated energy production to promote multiple myeloma cell proliferation. Cell Death Dis. 2022 Sep 26;13(9):822.

Cheng J, Dong Y, Ma J, Pan R, Liao Y, Kong X, Li X, Li S, Chen P, Wang L, Yu Y, Yuan Z. Microglial Calhm2 regulates neuroinflammation and contributes to Alzheimer’s disease pathology. Sci Adv. 2021 Aug 25;7(35):eabe3600.

Cheon SY, Kim EJ, Kim SY, Kim JM, Kam EH, Park JK, Koo BN. Apoptosis Signal-regulating Kinase 1 Silencing on Astroglial Inflammasomes in an Experimental Model of Ischemic Stroke. Neuroscience. 2018 Oct 15;390:218–230.

Chiarini A, Armato U, Gui L, Yin M, Chang S, Dal Prà I. Early divergent modulation of NLRP2’s and NLRP3’s inflammasome sensors vs. AIM2’s one by signals from Aβ·Calcium- sensing receptor complexes in human astrocytes. Brain Res. 2025 Jan 1;1846:149283.

Chiu TY, Lazar DC, Wang WW, Wozniak JM, Jadhav AM, Li W, Gazaniga N, Theofilopoulos AN, Teijaro JR, Parker CG. Chemoproteomic development of SLC15A4 inhibitors with anti-inflammatory activity. Nat Chem Biol. 2024 Aug;20(8):1000–1011.

Cossée M, Dürr A, Schmitt M, Dahl N, Trouillas P, Allinson P, Kostrzewa M, Nivelon- Chevallier A, Gustavson KH, Kohlschütter A, Müller U, Mandel JL, Brice A, Koenig M, Cavalcanti F, Tammaro A, De Michele G, Filla A, Cocozza S, Labuda M, Montermini L, Poirier J, Pandolfo M. Friedreich’s ataxia: point mutations and clinical presentation of compound heterozygotes. Ann Neurol. 1999 Feb;45(2):200–6.

Dagher M, Kleinman M, Ng A, Juncker D. Ensemble multicolour FRET model enables barcoding at extreme FRET levels. Nat Nanotechnol. 2018 Oct;13(10):925–932.

Dagher M, Ongo G, Robichaud N, Kong J, Rho W, Teahulos I, Tavakoli A, Bovaird S, Merjaneh S, Tan A, Edwardson K, Scheepers C, Ng A, Hajjar A, Sow B, Vrouvides M, Lee A, DeCorwin-Martin P, Rasool S, Huang J, Erps T, Coffin S, Rashidi NM, Han Y, Chandrasekaran SN, Miller L, Kost-Alimova M, Skepner A, Singh S, Carpenter AE, Munzar JD, Juncker D. nELISA: a high-throughput, high-plex platform enables quantitative profiling of the inflammatory secretome. Nat Methods. 2025 Nov;22(11):2375–2385.

de Alba E. Structure, interactions and self-assembly of ASC-dependent inflammasomes. Arch Biochem Biophys. 2019 Jul 30;670:15–31.

Della Valle I, Milani M, Rossi S, Turchi R, Tortolici F, Nesci V, Ferri A, Valle C, Lettieri-Barbato D, Aquilano K, Cozzolino M, Apolloni S, D’Ambrosi N. Loss of homeostatic functions in microglia from a murine model of Friedreich’s ataxia. Genes Dis. 2023 Nov 23;11(6):101178.

Deng Y, Wang D, Wang C, Guo K, Lei M, Huang Y, Tang L, Ding Y, Gao Y. Serum CHI3L1 levels correlate with disease activity in rheumatoid arthritis and reveal potential molecular mechanisms. Front Immunol. 2025 Dec 18;16:1729989.

Dinesh P, Rasool M. uPA/uPAR signaling in rheumatoid arthritis: Shedding light on its mechanism of action. Pharmacol Res. 2018 Aug;134:31–39.

Dobin A, Davis CA, Schlesinger F, Drenkow J, Zaleski C, Jha S, Batut P, Chaisson M, Gingeras TR. STAR: ultrafast universal RNA-seq aligner. Bioinformatics. 2013 Jan 1;29(1):15–21.

Dominic A, Le NT, Takahashi M. Loop Between NLRP3 Inflammasome and Reactive Oxygen Species. Antioxid Redox Signal. 2022 Apr;36(10-12):784–796.

Douvaras P, Buenaventura DF, Sun B, Lepack A, Baker E, Simpson E, Ebel M, Lallos G, LoSchiavo D, Stitt N, Adams N, McAuliffe C, Forton-Juarez A, Kosmyna B, Pereira E, Burnett B, Dilworth D, Fisher S, Wang J, Tonge P, Tomishima M, Paladini C, Wilkinson D, Soh CL, Srinivas M, Patsch C, Irion S. Ready-to-use iPSC-derived microglia progenitors for the treatment of CNS disease in mouse models of neuropathic mucopolysaccharidoses. Nat Commun. 2024 Sep 16;15(1):8132.

Ducza L, Gaál B. The Neglected Sibling: NLRP2 Inflammasome in the Nervous System. Aging Dis. 2024 May 7;15(3):1006–1028.

Farhadova S, Ghousein A, Charon F, Surcis C, Gomez-Velazques M, Roidor C, Di Michele F, Borensztein M, De Sario A, Esnault C, Noordermeer D, Moindrot B, Feil R. The long non-coding RNA Meg3 mediates imprinted gene expression during stem cell differentiation. Nucleic Acids Res. 2024 Jun 24;52(11):6183–6200.

Friker LL, Scheiblich H, Hochheiser IV, Brinkschulte R, Riedel D, Latz E, Geyer M, Heneka MT. β-Amyloid Clustering around ASC Fibrils Boosts Its Toxicity in Microglia. Cell Rep. 2020 Mar 17;30(11):3743–3754.e6.

Fu J, Wu H. Structural Mechanisms of NLRP3 Inflammasome Assembly and Activation. Annu Rev Immunol. 2023. Apr 26;41:301-316.

Galatro TF, Holtman IR, Lerario AM, Vainchtein ID, Brouwer N, Sola PR, Veras MM, Pereira TF, Leite REP, Möller T, Wes PD, Sogayar MC, Laman JD, den Dunnen W, Pasqualucci CA, Oba-Shinjo SM, Boddeke EWGM, Marie SKN, Eggen BJL. Transcriptomic analysis of purified human cortical microglia reveals age-associated changes. Nat Neurosci. 2017 Aug;20(8):1162–1171.

Gaudet RG, Zhu S, Halder A, Kim BH, Bradfield CJ, Huang S, Xu D, Mamiñska A, Nguyen TN, Lazarou M, Karatekin E, Gupta K, MacMicking JD. A human apolipoprotein L with detergent-like activity kills intracellular pathogens. Science. 2021 Jul 16;373(6552):eabf8113.

Guzmán-Guzmán IP, Ramírez-Vélez CI, Falfán-Valencia R, Navarro-Zarza JE, Gutiérrez-Pérez IA, Zaragoza-García O, Ramírez M, Castro-Alarcón N, Parra-Rojas I. *PADI2* polymorphisms are significantly associated with heumatoid Arthritis, autoantibodies serologic status and joint damage in women from southern mexico. Front Immunol. 2021 Aug 4;12:718246.

Haenseler W, Sansom SN, Buchrieser J, Newey SE, Moore CS, Nicholls FJ, Chintawar S, Schnell C, Antel JP, Allen ND, Cader MZ, Wade-Martins R, James WS, Cowley SA. A Highly Efficient Human Pluripotent Stem Cell Microglia Model Displays a Neuronal-Co-culture-Specific Expression Profile and Inflammatory Response. Stem Cell Reports. 2017 Jun 6;8(6):1727–1742.

Hara H, Tsuchiya K, Kawamura I, Fang R, Hernandez-Cuellar E, Shen Y, Mizuguchi J, Schweighoffer E, Tybulewicz V, Mitsuyama M. Phosphorylation of the adaptor ASC acts as a molecular switch that controls the formation of speck-like aggregates and inflammasome activity. Nat Immunol. 2013 Dec;14(12):1247–55.

Harris J, Lang T, Thomas JPW, Sukkar MB, Nabar NR, Kehrl JH. Autophagy and inflammasomes. Mol Immunol. 2017 Jun;86:10–15.

Healy, L. M., Yaqubi, M., Ludwin, S. & Antel, J. P. Species differences in immune-mediated CNS tissue injury and repair: A (neuro)inflammatory topic. Glia 68, 811–829 (2020).

Heidari A, Yazdanpanah N, Rezaei N. The role of Toll-like receptors and neuroinflammation in Parkinson’s disease. J Neuroinflammation. 2022 Jun 6;19(1):135.

Hickman SE, Kingery ND, Ohsumi TK, Borowsky ML, Wang LC, Means TK, El Khoury J. The microglial sensome revealed by direct RNA sequencing. Nat Neurosci. 2013 Dec;16(12):1896–905.

Hu G, Shen S, Zhu M. CXCL9 is a dual-role biomarker in colorectal cancer linked to mitophagy and modulated by ALKBH5. Mol Med Rep. 2025 Jul;32(1):188.

Imbault V, Dionisi C, Naeije G, Communi D, Pandolfo M. Cerebrospinal Fluid Proteomics in Friedreich Ataxia Reveals Markers of Neurodegeneration and Neuroinflammation. Front Neurosci. 2022 Jul 13;16:885313.

Ji DX, Witt KC, Kotov DI, Margolis SR, Louie A, Chevée V, Chen KJ, Gaidt MM, Dhaliwal HS, Lee AY, Nishimura SL, Zamboni DS, Kramnik I, Portnoy DA, Darwin KH, Vance RE. Role of the transcriptional regulator SP140 in resistance to bacterial infections via repression of type I interferons. Elife. 2021 Jun 21;10:e67290.

Jia C, Tan Y, Zhao M. The complement system and autoimmune diseases. Chronic Dis Transl Med. 2022 Apr 6;8(3):184–190.

Juliar BA, Stanaway IB, Sano F, Fu H, Smith KD, Akilesh S, Scales SJ, El Saghir J, Bhatraju PK, Liu E, Yang J, Lin J, Eddy S, Kretzler M, Zheng Y, Himmelfarb J, Harder JL, Freedman BS. Interferon-γ induces combined pyroptotic angiopathy and APOL1 expression in human kidney disease. Cell Rep. 2024 Jun 25;43(6):114310.

Khan W, Corben LA, Bilal H, Vivash L, Delatycki MB, Egan GF, Harding IH. Neuroinflammation in the Cerebellum and Brainstem in Friedreich Ataxia: An [18F]-FEMPA PET Study. Mov Disord. 2022 Jan;37(1):218–224.

Koeppen AH, Ramirez RL, Yu D, Collins SE, Qian J, Parsons PJ, Yang KX, Chen Z, Mazurkiewicz JE, Feustel PJ. Friedreich’s ataxia causes redistribution of iron, copper, and zinc in the dentate nucleus. Cerebellum. 2012 Dec;11(4):845–60.

Li B, Dewey CN. RSEM: accurate transcript quantification from RNA-Seq data with or without a reference genome. BMC Bioinformatics. 2011 Aug 4;12:323.

Li Y, Banerjee S, Wang Y, Goldstein SA, Dong B, Gaughan C, Silverman RH, Weiss SR. Activation of RNase L is dependent on OAS3 expression during infection with diverse human viruses. Proc Natl Acad Sci U S A. 2016 Feb 23;113(8):2241–6.

Li N, Du J, Yang Y, Zhao T, Wu D, Peng F, Wang D, Kong L, Zhou W, Hao A. Microglial PCGF1 alleviates neuroinflammation associated depressive behavior in adolescent mice. Mol Psychiatry. 2025 Mar;30(3):914–926.

Liu Q, Zhang D, Hu D, Zhou X, Zhou Y. The role of mitochondria in NLRP3 inflammasome activation. Mol Immunol. 2018 Nov;103:115–124.

Llorens JV, Soriano S, Calap-Quintana P, Gonzalez-Cabo P, Moltó MD. The Role of Iron in Friedreich’s Ataxia: Insights From Studies in Human Tissues and Cellular and Animal Models. Front Neurosci. 2019 Feb 18;13:75.

López-Haber C, Netting DJ, Hutchins Z, Ma X, Hamilton KE, Mantegazza AR. The phagosomal solute transporter SLC15A4 promotes inflammasome activity via mTORC1 signaling and autophagy restraint in dendritic cells. EMBO J. 2022 Oct 17;41(20):e111161.

Losanto J, Langjahr P, Barrios G, Paats A, Acosta de Hetter ME, de Guillén I, Duarte M, Acosta-Colman I, Cervera R. Relationship between serum lipopolysaccharide binding protein levels, disease activity, and clinical characteristics in Paraguayan patients with systemic lupus erythematosus. Lupus. 2021 Nov;30(13):2089–2094.

Love MI, Huber W, Anders S. Moderated estimation of fold change and dispersion for RNA-seq data with DESeq2. Genome Biol. 2014;15(12):550.

Lu JQ, Wang HQ, Fang MR, Ye XM, Song KY. Autophagy-NLRP3 Inflammasome Crosstalk in Microglia: A Therapeutic Target for Multiple Sclerosis. Inflammation. 2026 Jan 3;49(1):23.

Ma H, Yin Q, Li H, Guo M. Differential complement pathways and components in the clinical spectrum of systemic lupus erythematosus. Arthritis Res Ther. 2025 Dec 26;28(1):23.

Ma SQ, Wang L, Liu X, Yue YN, Wang SN, Feng Y, Xin S, Zhu P, Li FS, Yin SM. PLAUR Exacerbates Neuroinflammation in Diabetic Ischemic Stroke by Driving Neutrophil-Mediated Blood-Brain Barrier Disruption and Reprogramming Microglial Metabolism. Neuromolecular Med. 2026 May 26;28(1):28.

Makoni NJ, Nichols MR. The intricate biophysical puzzle of caspase-1 activation. Arch Biochem Biophys. 2021 Mar 15;699:108753.

Masuda T, Sankowski R, Staszewski O, Böttcher C, Amann L, Sagar, Scheiwe C, Nessler S, Kunz P, van Loo G, Coenen VA, Reinacher PC, Michel A, Sure U, Gold R, Grün D, Priller J, Stadelmann C, Prinz M. Spatial and temporal heterogeneity of mouse and human microglia at single-cell resolution. Nature. 2019 Feb;566(7744):388–392.

McMurray JC, Schornack BJ, Weskamp AL, Park KJ, Pollock JD, Day WG, Brockshus AT, Beakes DE, Schwartz DJ, Mikita CP, Pittman LM. Immunodeficiency: Complement disorders. Allergy Asthma Proc. 2024 Sep 1;45(5):305–309.

Meng J, Ding T, Chen Y, Long T, Xu Q, Lian W, Liu W. LncRNA-Meg3 promotes Nlrp3- mediated microglial inflammation by targeting miR-7a-5p. Int Immunopharmacol. 2021 Jan;90:107141.

Meng L, Wang J, Chen H, Zhu J, Kong F, Chen G, Dong R, Zheng S. LncRNA MEG9 Promotes Inflammation and Liver Fibrosis Through S100A9 in Biliary Atresia. J Pediatr Surg. 2025 Feb;60(2):161633.

Metkar SS, Menaa C, Pardo J, Wang B, Wallich R, Freudenberg M, Kim S, Raja SM, Shi L, Simon MM, Froelich CJ. Human and mouse granzyme A induce a proinflammatory cytokine response. Immunity. 2008 Nov 14;29(5):720–33.

Minkiewicz J, de Rivero Vaccari JP, Keane RW. Human astrocytes express a novel NLRP2 inflammasome. Glia. 2013 Jul;61(7):1113–21.

Mishra N, Schwerdtner L, Sams K, Mondal S, Ahmad F, Schmidt RE, Coonrod SA, Thompson PR, Lerch MM, Bossaller L. Cutting Edge: Protein Arginine Deiminase 2 and 4 Regulate NLRP3 Inflammasome-Dependent IL-1β Maturation and ASC Speck Formation in Macrophages. J Immunol. 2019 Aug 15;203(4):795–800.

Paik S, Kim JK, Shin HJ, Park EJ, Kim IS, Jo EK. Updated insights into the molecular networks for NLRP3 inflammasome activation. Cell Mol Immunol. 2025 Jun;22(6):563–596.

Palaparti A, Baratz A, Stifani S. The Groucho/transducin-like enhancer of split transcriptional repressors interact with the genetically defined amino-terminal silencing domain of histone H3. J Biol Chem. 1997 Oct 17;272(42):26604–10.

Pernaci C, Johnson A, Gillette S, Warden AS, McCormick C, Weiser-Novak S, Ramirez G, Broersma EH, Mishra P, Sivakumar A, Cherqui S, Coufal NG. Microgliopathy as a primary mediator of neuronal death in models of Friedreich’s Ataxia. Nat Commun. 2025 Nov 29.

Prather ER, Gavrilin MA, Wewers MD. The central inflammasome adaptor protein ASC activates the inflammasome after transition from a soluble to an insoluble state. J Biol Chem. 2022 Jun;298(6):102024.

Reetz K, Lischewski SA, Dogan I, Didszun C, Pishnamaz M, Konrad K, Marx-Schütt K, Farmer J, Lynch DR, Corben LA, Pandolfo M, Schulz JB; FACROSS study group. Friedreich’s ataxia-a rare multisystem disease. Lancet Neurol. 2025 Jul;24(7):614–624.

Rimann I, Gonzalez-Quintial R, Baccala R, Kiosses WB, Teijaro JR, Parker CG, Li X, Beutler B, Kono DH, Theofilopoulos AN. The solute carrier SLC15A4 is required for optimal trafficking of nucleic acid-sensing TLRs and ligands to endolysosomes. Proc Natl Acad Sci U S A. 2022 Apr 5;119(14):e2200544119.

Ristow M, Pfister MF, Yee AJ, Schubert M, Michael L, Zhang CY, Ueki K, Michael MD 2nd, Lowell BB, Kahn CR. Frataxin activates mitochondrial energy conversion and oxidative phosphorylation. Proc Natl Acad Sci U S A. 2000 Oct 24;97(22):12239–43.

Ritacco DA, Shahnawaz H, Oduguwa A, Hawk J, Vizcaino B, Farber DL, Gaudet RG. The human antibacterial factor APOL3 couples lysosomal damage to mitochondrial DNA efflux and type I IFN induction. Mol Cell. 2026 Mar 19;86(6):1116–1133.e8.

Sandstrom A, Vance RE. Defusing inflammasomes. J Exp Med. 2018 Mar 5;215(3):723–724.

Sciarretta F, Zaccaria F, Ninni A, Ceci V, Turchi R, Apolloni S, Milani M, Della Valle I, Tiberi M, Chiurchiù V, D’Ambrosi N, Pedretti S, Mitro N, Volontè C, Amadio S, Aquilano K, Lettieri-Barbato D. Frataxin deficiency shifts metabolism to promote reactive microglia via glucose catabolism. Life Sci Alliance. 2024 Apr 17;7(7):e202402609.

Sharif M, Anvar Z, Chakchouk I, El-Dessouky SH, Zemet R, Kao EC, Sharaf-Eldin WE, Wan YW, Liu Z, Liu P, Jochum M, Van den Veyver IB. Loss of the maternal effect gene NLRP2 impairs embryonic and extra-embryonic development, revealing a novel genetic cause of congenital anomalies†. Biol Reprod. 2026 Apr 13;114(4):1469–1485.

Sharma BR, Kanneganti TD. NLRP3 inflammasome in cancer and metabolic diseases. Nat Immunol. 2021 May;22(5):550–559.

Shen Y, McMackin MZ, Shan Y, Raetz A, David S, Cortopassi G. Frataxin Deficiency Promotes Excess Microglial DNA Damage and Inflammation that Is Rescued by PJ34. PLoS One. 2016 Mar 8;11(3):e0151026.

Shi Q, Gutierrez RA, Bhat MA. Microglia, Trem2, and Neurodegeneration. Neuroscientist. 2025 Apr;31(2):159–176.

Shiau CE, Monk KR, Joo W, Talbot WS. An anti-inflammatory NOD-like receptor is required for microglia development. Cell Rep. 2013 Dec 12;5(5):1342–52.

Sogorb-Esteve A, Swift IJ, Woollacott IOC, Warren JD, Zetterberg H, Rohrer JD. Differential chemokine alteration in the variants of primary progressive aphasia-a role for neuroinflammation. J Neuroinflammation. 2021 Oct 3;18(1):224.

Sun Y, Xie X, Zou X, Zhou F. Neuroinflammatory chemokine networks in transgenic models of Alzheimer’s disease: A comparative multi-compartmental analysis. Hum Exp Toxicol. 2025 Jan-Dec;44:9603271251348723.

Tang YM, Pulimood NS, Stifani S. Comparing the Characteristics of Microglia Preparations Generated Using Different Human iPSC-Based Differentiation Methods to Model Neurodegenerative Diseases. ASN Neuro. 2022 Jan-Dec;14:17590914221145105.

Terashima M, Ishimura A, Wanna-Udom S, Suzuki T. *MEG8* long noncoding RNA contributes to epigenetic progression of the epithelial-mesenchymal transition of lung and pancreatic cancer cells. J Biol Chem. 2018 Nov 23;293(47):18016–18030.

Thiry L, Pulimood NS, Tang YM, Stifani S. Dysregulated Expression of Inflammasome and Extracellular Matrix Genes in *C9orf72*-ALS/FTD Microglia. ASN Neuro. 2025;17(1):2542998.

Vajjhala PR, Mirams RE, Hill JM. Multiple binding sites on the pyrin domain of ASC protein allow self-association and interaction with NLRP3 protein. J Biol Chem. 2012 Dec 7;287(50):41732–43.

Vervliet T, Loncke J, Sever M, Ahuja K, Van den Haute C, Luyten T, Stutzmann GE, Verfaillie C, Tomašič T, Bultynck G. Inactive ryanodine receptors sustain lysosomal availability for autophagy by promoting ER-lysosomal contact site formation. Nat Commun. 2026 Jan 15;17(1):1293.

Vicente-Acosta A, Herranz-Martín S, Pazos MR, Galán-Cruz J, Amores M, Loria F, Díaz-Nido J. Glial cell activation precedes neurodegeneration in the cerebellar cortex of the YG8-800 murine model of Friedreich ataxia. Neurobiol Dis. 2024 Oct 1;200:106631.

Wang H, Brown J, Gao S, Liang S, Jotwani R, Zhou H, Suttles J, Scott DA, Lamont RJ. The role of JAK-3 in regulating TLR-mediated inflammatory cytokine production in innate immune cells. J Immunol. 2013 Aug 1;191(3):1164–74.

Wei C, Liu J, Wu B, Shen T, Fan J, Lin Y, Li K, Guo Y, Shang Y, Zhou B, Xie H. Blockage of CCL3 with neutralizing antibody reduces neuroinflammation and reverses Alzheimer disease phenotypes. Brain Behav Immun. 2025 Aug;128:400–415.

Wu J, Raman A, Coffey NJ, Sheng X, Wahba J, Seasock MJ, Ma Z, Beckerman P, Laczkó D, Palmer MB, Kopp JB, Kuo JJ, Pullen SS, Boustany-Kari CM, Linkermann A, Susztak K. The key role of NLRP3 and STING in APOL1-associated podocytopathy. J Clin Invest. 2021 Oct 15;131(20):e136329.

Xu H, Ma H, Zha L, Li Q, Pan H, Zhang L. Engineered exosomes transporting the lncRNA, SVIL-AS1, inhibit the progression of lung cancer via targeting miR-21-5p. Am J Cancer Res. 2024a Jul 15;14(7):3335-3347.

Xu J, Gu J, Pei W, Zhang Y, Wang L, Gao J. The role of lysosomal membrane proteins in autophagy and related diseases. FEBS J. 2024b Sep;291(17):3762–3785.

Ye Y, Wang M, Wang G, Mai Z, Zhou B, Han Y, Zhuang J, Xia W. lncRNA miR4458HG modulates hepatocellular carcinoma progression by activating m6A-dependent glycolysis and promoting the polarization of tumor-associated macrophages. Cell Mol Life Sci. 2023 Mar 18;80(4):99.

Yu TG, Cha JS, Kim G, Sohn YK, Yoo Y, Kim U, Song JJ, Cho HS, Kim HS. Oligomeric states of ASC specks regulate inflammatory responses by inflammasome in the extracellular space. Cell Death Discov. 2023 Apr 29;9(1):142.

Zahl S, Skauli N, Stahl K, Prydz A, Frey MM, Dissen E, Ottersen OP, Amiry-Moghaddam M. Aquaporin-9 in the Brain Inflammatory Response: Evidence from Mice Injected with the Parkinsonogenic Toxin MPP. Biomolecules. 2023 Mar 24;13(4):588.

Zang X, Wang J, Xia Y, Li J, Chen L, Gu Y, Shen X. LncRNA MEG3 promotes the sensitivity of bortezomib by inhibiting autophagy in multiple myeloma. Leuk Res. 2022 Dec;123:106967.

Zhang L, Mo J, Swanson KV, Wen H, Petrucelli A, Gregory SM, Zhang Z, Schneider M, Jiang Y, Fitzgerald KA, Ouyang S, Liu ZJ, Damania B, Shu HB, Duncan JA, Ting JP. NLRC3, a member of the NLR family of proteins, is a negative regulator of innate immune signaling induced by the DNA sensor STING. Immunity. 2014 Mar 20;40(3):329–41.

Zhang T, Shu Q, Zhu H, Wang M, Yang N, Zhang H, Ge W. Serum proteomics analysis of biomarkers for evaluating clinical response to MTX/IGU therapy in early rheumatoid arthritis. Mol Immunol. 2023 Jan;153:119–125.

Zhang T, Xing F, Qu M, Yang Z, Liu Y, Yao Y, Xing N. NLRP2 in health and disease. Immunology. 2024 Feb;171(2):170–180.

Zhu F, Li S, Gu Q, Xie N, Wu Y. APOL1 Induces Pyroptosis of Fibroblasts Through NLRP3/Caspase-1/GSDMD Signaling Pathway in Ulcerative Colitis. J Inflamm Res. 2023 Dec 27;16:6385–6396.

